# Arabinogalactans provide the molecular basis of ice nucleation by plant pollen and spores

**DOI:** 10.64898/2026.11.25.769183

**Authors:** Nina L. H. Kinney, Emma Cooper, Fabienne Bachtiger, Akalabya Bissoyi, Douglas E. Soutar, Martin I. Daily, Caroline I. Biggs, Ieva Lelėnaitė, Charlotte Fletcher, Collette S. Guy, Alexander N. Baker, J. Paul Knox, William G. T. Willats, Daniel Ballesteros, Matthew I. Gibson, Gabriele C. Sosso, Thomas F. Whale

**Affiliations:** School of Earth, Environment and Sustainability, University of Leeds, Leeds LS2 9JT, UK; Department of Chemistry, University of Warwick, Coventry CV4 7AL, UK; Manchester Institute of Biotechnology, University of Manchester, Manchester M1 7DN, UK; School of Natural and Environmental Sciences, Newcastle University, Newcastle upon Tyne NE1 7RU, UK; School of Life Sciences, University of Warwick, Coventry CV4 7AL, UK; Warwick Medical School, University of Warwick, Coventry CV4 7AL, UK; Centre for Plant Sciences, Faculty of Biological Sciences, University of Leeds, Leeds LS2 9JT, UK; Department of Botany and Geology, University of Valencia, 46100 Burjassot, Valencia, Spain; Trait Diversity and Function, Royal Botanic Gardens, Kew, Ardingly, West Sussex RH17 6TN, UK; Department of Chemistry, University of Manchester, Manchester M13 9PL, UK

## Abstract

Biological ice nucleators are thought to influence cloud ice formation, affecting weather and climate. Plants are a major source of atmospheric aerosol, yet the identity of the ice-nucleating molecules they produce remains unclear. Droplet freezing experiments show that pollen, fern spores and moss spores carry similar soluble ice-nucleating macromolecules, indicating that such material occurs across land plants. Selective binding experiments and infrared spectroscopy identify these molecules as Type II arabinogalactans or closely related polysaccharides. Size fractionation, molecular simulations and classical nucleation theory indicate that these ubiquitous, structurally variable molecules nucleate ice only in sufficiently large, suitably organised assemblies. We conclude that plant ice nucleation is an incidental property of arabinogalactan assemblies, rather than necessarily an evolved adaptation, explaining its widespread but highly variable occurrence.

## Main Text

Ice nucleation from supercooled water is a critical process in many environmental and technological settings, yet its molecular basis remains poorly understood. In the absence of ice-nucleating particles (INPs), liquid water can supercool as low as –38 °C before homogeneous nucleation occurs (*1*). By contrast, some INPs can induce heterogeneous nucleation at temperatures close to the melting point of water. Heterogeneous ice nucleation by INPs plays a key role in determining radiative and precipitative properties of supercooled clouds, thereby influencing weather and climate (*2, 3*). The climatic influence of INPs remains uncertain because their atmospheric abundance and effects are poorly constrained (*4, 5*).

Pollen grains and fragments from anemophilous (wind-pollinating) plants are an important component of atmospheric aerosol. Global pollen emissions are estimated at 84.5 × 10^9^ kg year^-1^ (*6*) and sub-pollen particle (SPP) concentrations are estimated to be between four and six orders of magnitude greater than pollen concentrations (*7*). Pollen from many plant species nucleates ice effectively (*7–17*). Ground and satellite-based observations indicate that pollen emissions enhance cloud glaciation (*18*) while modelling studies indicate that pollen ice nucleation can substantially affect cloud properties, especially when pollen rupture and SPP emission are included (*7, 19–21*).

Pummer et al. (*10*) showed that the ice-active agents produced by some pollens are water-soluble ice-nucleating macromolecules (INMs) that separate readily from pollen grains. Despite considerable subsequent research, their molecular identity remains unclear. There is evidence these INMs are polysaccharides (*11, 17*), unlike the proteinaceous INMs produced by bacteria and fungi (*22, 23*). However, protein-compatible fluorescence and responses to phenol extraction suggest that a protein component may contribute to the activity of at least some pollen INMs (*14, 17, 24, 25*).

Here, we show that fern and moss spores release soluble INMs of the same class as those produced by seed-plant pollen, extending their known occurrence across the land-plant phylogeny and tracing the origin of these molecules to at least 494 million years ago (*26*). We identify Type II arabinogalactans, ubiquitous and structurally diverse plant cell-wall polysaccharides, as the likely molecules responsible for this activity. Binding and size-fractionation experiments suggest that INA arises when ubiquitous plant macromolecules happen to form sufficiently large and suitably organised assemblies, potentially explaining the widespread but variable occurrence of INA across land plants without requiring the evolution of a specialised function.

### Fern and moss spores produce ice-nucleating macromolecules

Our recent study of ice nucleation by pollen INMs across the seed plant phylogeny demonstrated highly variable ice-nucleating activity (INA) between species (*15*). We found no association between INA and the plant or pollen characteristics examined, suggesting that pollen INMs are widespread and that their INA is likely incidental rather than an adaptive trait (*15*).

The widespread occurrence of pollen INMs led us to ask whether similar material is also produced by more distantly related land plants. Before the evolution of pollen-producing seed plants in the late Devonian (*27*), land-plant reproduction and dispersal relied on spores, as it still does in extant ferns and bryophytes (hornworts, liverworts and mosses). Despite their functional differences, pollen and spores are considered homologous structures, and much of the molecular machinery underlying their development is conserved (*28*).

We found that fern spores release water-soluble INMs spanning a wide range of activities. In some samples, soluble material associated with fern spores nucleated ice at temperatures approaching -5 °C. Spore washing water (SWW) was prepared by suspending spores in purified water and removing the spores and insoluble material by filtration (see Fig. S1 for schematic and Table S1 for sample collection details). Ice nucleation temperatures were measured using a microlitre droplet freezing experiment (µ-DFE) (*29*). Results for selected samples are shown in Fig. 1A and 1B, with the complete fern SWW dataset in Figs. S2 and S3. Unfiltered spore suspensions and filtered SWW showed similar INA, indicating that the active material separates readily from the spores, as reported previously for pollen washing water (PWW) (*10, 15, 30*).

**Fig. 1.**
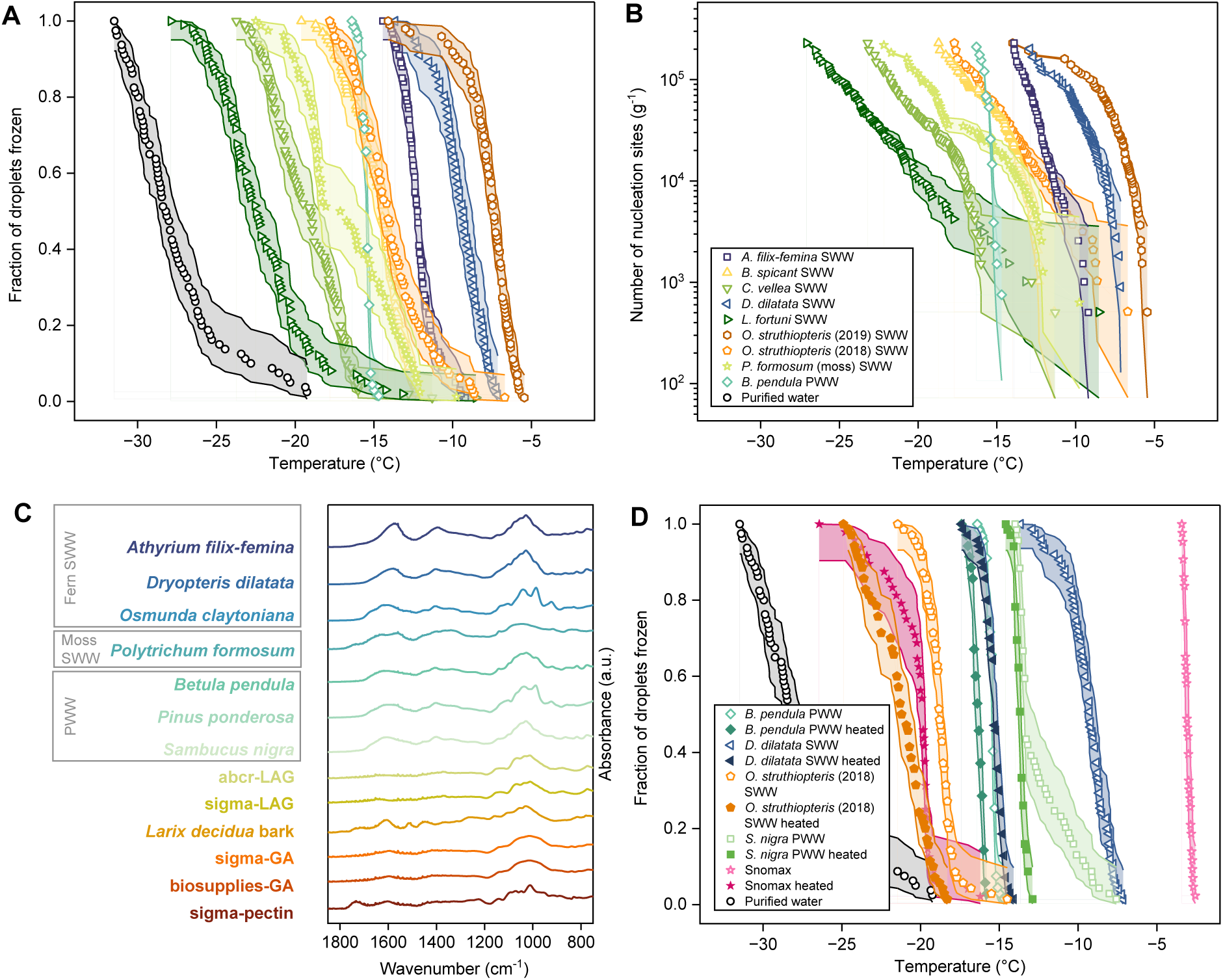
Fern and moss spores produce ice-nucleating macromolecules similar to those produced by angiosperm and gymnosperm pollens. (A) Fraction of 1 µL droplets frozen as a function of temperature for 2 wt% SWW from selected fern species and the moss Polytrichum formosum. A 2 wt% Betula pendula PWW sample and purified water are included for comparison. Multiple collections of Onoclea struthiopteris are identified by year. (B) Cumulative number of nucleation sites per unit dry mass, nm(T), as a function of temperature for the same fern and moss spore samples compared with Betula pendula PWW. (C) Normalised FT-IR absorption spectra of dried PWW and SWW residues, ground Larix decidua bark, commercial larch arabinogalactans, gum Arabic (GA) and pectin in the fingerprint region, 750–1850 cm⁻¹. Suppliers are indicated where relevant. (D) Fraction of droplets frozen for selected SWW and PWW samples and 0.05 wt% Snomax® before and after heating.

Spores from the bank haircap moss (*Polytrichum formosum*) also released soluble INMs (see Table S2 for collection details). *P. formosum* SWW nucleated ice at temperatures above –12 °C (Fig. 1A) and removing the spores did not substantially alter its activity (Fig. S4). Thus, soluble INMs are produced by mosses as well as ferns and seed plants.

FT-IR absorption spectra of freeze-dried fern and moss SWW residues closely resembled those of PWW residues (Fig. 1C). As previously reported for PWW (*17*), these spectra are consistent with samples dominated by polysaccharides, although individual compounds cannot be assigned confidently in complex natural extracts (*31*). Further analysis of the spectra may be found in the supplementary text. Together, the freezing and spectroscopic measurements indicate that pollen, fern spores and moss spores release soluble, predominantly polysaccharide INMs with similar properties.

We next used heat treatment (denaturing) to examine whether proteins contribute to the activity of these INMs (Fig. 1D). Heat treatment is known to denature and decrease INA from protein-containing INMs (almost eliminating the activity of the proteinaceous ice nucleator Snomax® (*32*)) and was hence applied to these samples. By contrast, *Betula pendula* PWW was largely heat-resistant, consistent with previous measurements, although other evidence suggests that its soluble extract contains a proteinaceous component (*32*). Heating substantially reduced the activity of the highly active *Dryopteris dilatata* SWW, lowering its median freezing temperature by 6 °C, and also reduced the activity of *Onoclea struthiopteris* SWW (Fig. 1D). *Sambucus nigra* PWW similarly lost a small population of freezing events between –13 and –7.5 °C as did the moss *Ceratodon purpureus* SWW between –15 and –6°C (Fig. S4B). After heating, these samples all retained steep freezing curves, with most events occurring between –13 and –16 °C.

These results suggest that heat-resistant polysaccharides account for much of the INA of pollen and spore extracts, while a heat-sensitive component contributes to the warmer-temperature activity of some samples. One possibility is that an associated protein promotes the formation or organisation of larger, more active polysaccharide assemblies, although heat treatment alone cannot establish this mechanism.

Together, our results show that moss and fern spores release soluble INMs resembling those produced by angiosperm and gymnosperm pollen. Their occurrence across these divergent lineages connects plant INA to a class of macromolecules distributed broadly across the land-plant phylogeny, originating before the divergence of vascular plants from bryophytes during the Cambrian Period (*26*) as illustrated in Fig. 2. Variation among fern SWWs, like that previously observed among PWWs (*15*), is consistent with ice nucleation being an incidental and highly variable property of these molecules.

**Fig. 2.**
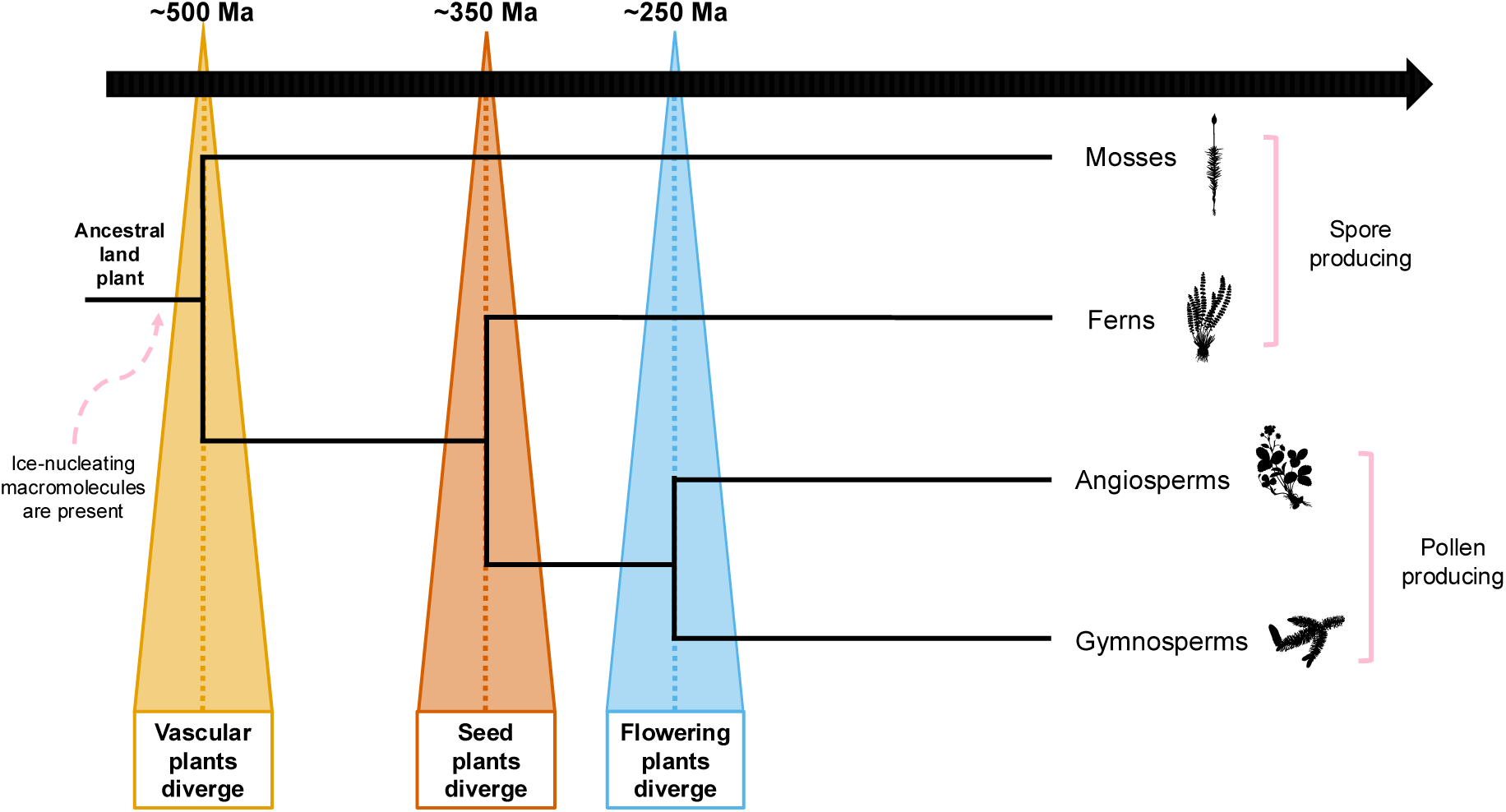
A shared class of soluble macromolecules nucleates ice across land plants. Phylogenetic tree showing relationships between extant land-plant groups found to release INMs, with a timeline of major events in plant evolution based on published estimates (*26, 33–35*).

### Arabinogalactans as candidate plant INMs

The similar properties of pollen and spore INMs suggested that their activity might arise from a molecular component shared across land plants. We therefore examined the soluble glycans in PWW and SWW using high-throughput microarray antigen profiling (Fig. S5). Several glycan epitopes were detected in PWW, including those associated with arabinan, homogalacturonan, xyloglucan, extensin and arabinogalactan protein (AGP). No epitopes were detected in the SWW samples. This negative result does not establish that glycans are absent from SWW: the antibodies recognise a restricted set of epitopes, and the concentration of individual glycans may have been below the detection limit. Consistent with this limitation, some AGPs from early-diverging vascular plants do not bind antibodies used in the array (*36*).

The combined evidence suggested that the plant INMs are soluble glycans that may, in some samples, be associated with protein. We therefore hypothesised that they are Type II arabinogalactans (AGs), which commonly occur attached to protein backbones asarabinogalactan proteins (AGPs). This could reconcile evidence that pollen INMs are predominantly polysaccharide with indications that a protein component contributes to the activity of some extracts (*14, 25, 37, 38*).

Type II AGs and AGPs occur throughout land plants, including mosses, ferns and seed plants (*36, 39–44*). They are highly branched, structurally variable and, unusually among plant cell-wall polysaccharides, soluble (*40, 42, 45*). These properties make Type II AGs plausible candidates for the widespread, soluble and variably active INMs released from pollen and spores.

### Plant arabinogalactans nucleate ice

To test whether arabinogalactans could account for the activity of plant INMs, we examined two commercial larch arabinogalactan preparations (abcr-LAG and Sigma-LAG), an AGP-containing gum arabic sample, pectin and a mycobacterial arabinogalactan (which has a distinct structure from plant AG). The FT-IR spectra of the plant-derived samples closely resembled those of the PWW and SWW residues (Fig. 1C), although these spectra are not diagnostic of AG.

All plant-derived polysaccharide samples nucleated ice above the instrument background at 1 wt% (Fig. 3A). Of these, abcr-LAG was the most active, producing a steep fraction-frozen curve with a median freezing temperature of –14 °C, comparable to many PWW and SWW samples. This steep transition, in which most droplets froze over a narrow temperature interval, is also characteristic of many PWW and SWW samples. Although not diagnostic of molecular identity, this similarity in freezing behaviour is consistent with related INMs being present in commercial LAG and natural plant extracts. By contrast, the mycobacterial (furanose form) arabinogalactan, which has consistent repeating structure, in contrast to plant AG (*46*), did not nucleate ice above the instrument background.

**Fig. 3.**
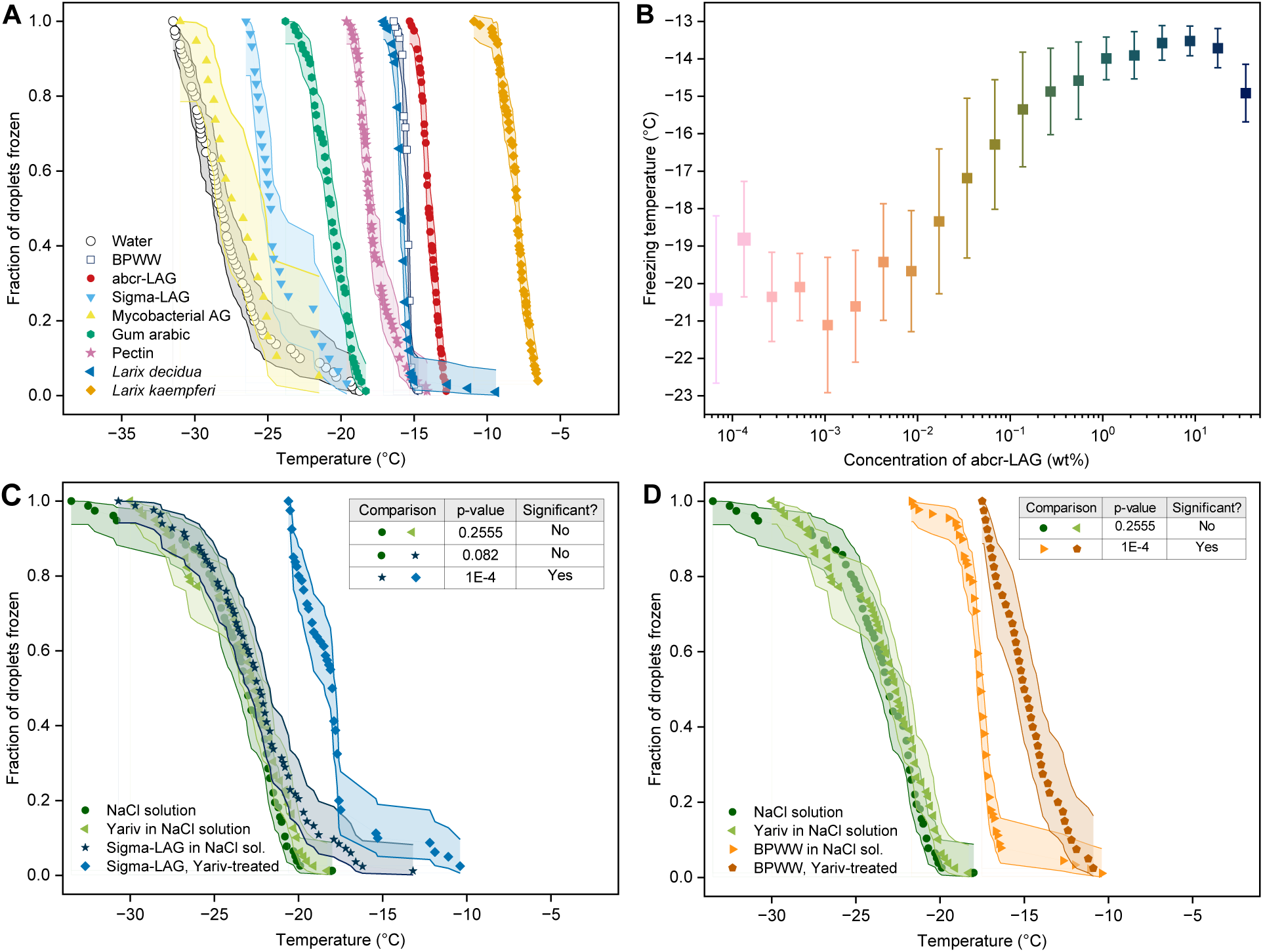
Arabinogalactan samples nucleate ice effectively. (A) Fraction of 1 µL droplets frozen as a function of temperature for 1 wt% solutions of plant-derived arabinogalactans and related polysaccharides. A 2 wt% *Betula pendula* pollen washing water (BPWW) sample and the purified-water instrument background are included for comparison. (B) Mean freezing temperature of 1 µL abcr-LAG droplets as a function of concentration. Whiskers show the highest and lowest measured droplet freezing temperatures. (C, D) Fraction of droplets frozen for 0.01 wt% Sigma-LAG and 0.01 wt% BPWW, respectively, in 0.15 M NaCl before and after addition of 0.5 mg mL⁻¹ β-glucosyl Yariv reagent. P-values were calculated using the Integrated Squared Difference test (*48*).

The INA of abcr-LAG depended non-monotonically on concentration (Fig. 3B). Ice nucleation temperatures increased with concentration up to 8.75 wt%, but decreased at higher concentrations, where melting-point depression increasingly affected the measured freezing temperatures. Thus, the highest measured freezing temperature reflects the competing effects of increasing nucleator abundance and increasing solute concentration. Ice recrystallisation and dynamic ice shaping measurements demonstrated that plant AGs bind to ice (Fig. S6, S7) a property that may be related to their ice-nucleating activity.

Filtered extracts of *Larix decidua* and *Larix kaempferi* bark were also strongly ice-active, with some droplets freezing at temperatures as high as –6 °C (Fig. 3A). FT-IR spectra of their freeze-dried residues exhibit the same dominant polysaccharide absorbance peaks as the commercial LAG samples, providing further evidence that plant tissues rich in AG are strongly ice active.

We next examined whether the active material contained the β-1,3-galactan backbone characteristic of Type II AG. Yariv phenylglycosides bind selectively to sufficiently long β-1,3-galactan chains and have therefore been widely used to identify Type II AGs and AGPs (*47*). Treating 0.01 wt% Sigma-LAG with β-glucosyl Yariv reagent increased its median ice nucleation temperature by 4.2 °C (Fig. 3C; *P* < 1 × 10^-4^), whereas Yariv reagent alone did not nucleate ice above the corresponding solution background (Fig. S8A). *Betula pendula* PWW showed a similar, although smaller, increase in activity following Yariv treatment (Fig. 3D; *P* < 1 × 10^-4^). The INA of the more active abcr-LAG was not significantly changed by Yariv treatment (Fig. S8D; *P* = 0.4381).

These results link the ice-nucleating material in Sigma-LAG and *Betula pendula* PWW to Type II AG. The increase in activity following Yariv treatment also indicates that β-1,3-galactan-mediated association can enhance ice nucleation. The absence of a corresponding response in abcr-LAG suggests that Yariv binding does not have a uniform effect across structurally heterogeneous AG samples, particularly where the untreated material is already strongly ice-active. Together with the distinctive steepness of the abcr-LAG freezing curve and the activity of larch bark extracts, these results provide converging evidence that Type II AG contributes to the soluble INMs released by plant material. Given the considerable structural diversity of plant AGs, INA may arise from several distinct AG structures and could extend to related polysaccharides with similar structural features.

### Molecular size and supramolecular organisation control arabinogalactan ice nucleation

To investigate how AG might promote ice nucleation, we used molecular dynamics simulations to test whether an isolated model LAG molecule reproducibly enhanced local ice-like ordering of water. Across five independent simulations, pre-critical ice-like clusters were not consistently enriched at the LAG–water interface, and measures of interfacial water ordering showed only marginal effects (Figs. S15 –S18). The simulations therefore provide no evidence that the ice-nucleating ability of LAG arises from the local interaction of an individual LAG molecule with water. However, the molecular sizes and timescales accessible to these simulations are much smaller than those implicated by the fractionation experiments. The structures responsible for experimental INA may instead arise from larger-scale association and organisation that are not represented by the model.

Similarly, the substantial variation in INA among AG samples indicates that chemical identity alone does not determine their ice-nucleating ability. Characterisation of abcr-LAG and Sigma-LAG using aqueous size exclusion chromatography (SEC) and nuclear magnetic resonance (NMR), detailed in the SI, revealed a small population of higher molecular weight species in the abcr-LAG sample (Figs. S9-S13). Previous studies of *Betula pendula* PWW found that its most active INMs occur in high-molecular-mass fractions, whereas smaller fractions nucleate ice at lower temperatures (*17, 30, 38*). We observed similar size-dependent activity in *Carpinus betulus* PWW (see Fig. S14 for fraction frozen curves and corresponding FT-IR spectra) and in size-fractionated Sigma-LAG (Fig. 4A). The most active Sigma-LAG material occurred in the fraction centred at approximately 100 kDa, while both smaller and larger fractions nucleated ice at lower temperatures. Thus, the activity of LAG does not increase monotonically with molecular size but is associated with particular high-molecular-mass fractions.

**Fig. 4.**
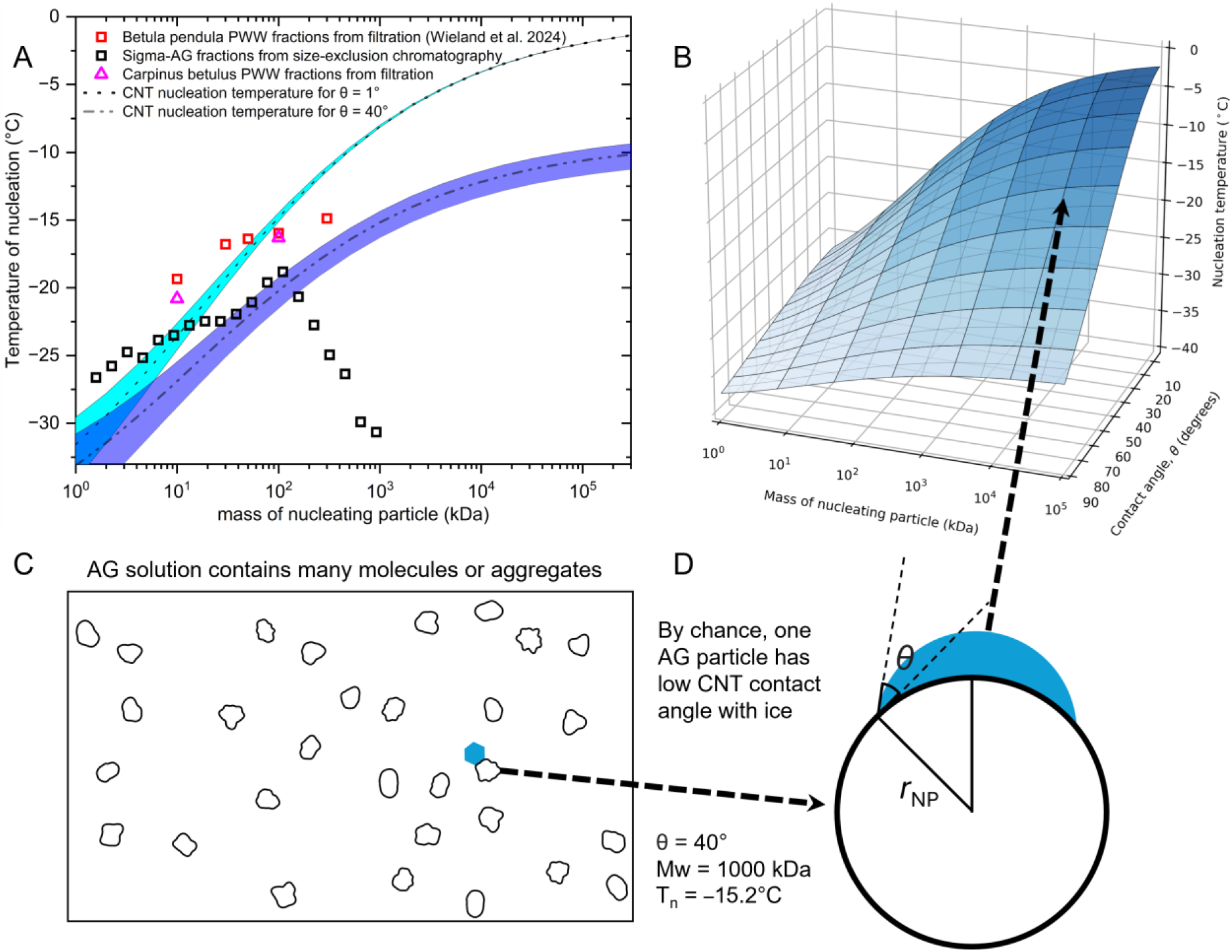
Molecular size and supramolecular organisation constrain arabinogalactan ice nucleation. **(A)** Median ice-nucleation temperatures of size-fractionated *Betula pendula* and *Carpinus betulus* pollen washing water (PWW) and larch arabinogalactan (LAG), plotted against estimated molecular mass. Dashed lines show classical nucleation theory (CNT) predictions for spherical nucleators with effective contact angles, θ, of 1° and 40°; shading reflects uncertainty in molecular mass. (B) CNT-predicted nucleation temperature as a function of nucleator mass and effective contact angle. Warm nucleation requires sufficient size and favourable interfacial properties. (C) A solution may contain many chemically similar arabinogalactan particles that differ in size and structure. (D) Only sufficiently large and favourably organised particles nucleate ice at warm temperatures, producing rare, highly active “chance nucleators”.

Molecular size places a physical constraint on heterogeneous ice nucleation because an active surface must be large enough to accommodate a critical ice nucleus. We examined this constraint using classical nucleation theory (CNT) for spherical ice clusters forming on spherical nucleators, following Fletcher (*49*). A discussion of the theory may be found in the supplement. Within this model, the nucleation temperature depends on both nucleator size and the effective contact angle between the nucleator and ice. Warm-temperature nucleation requires a sufficiently large nucleator together with favourable interfacial properties (Fig. 4 B, D). For example, the model predicts that a particle of 100 kDa or smaller cannot nucleate ice above approximately –14.7 °C, even at an effective contact angle of 1°. Although the assumed geometry is highly simplified, the calculation shows that molecular association is necessary for CNT to explain the warmest freezing events observed for LAG and plant extracts.

Size alone, however, cannot explain our measurements. Sigma-LAG fractions of similar nominal molecular mass differed in activity from pollen INMs, while the largest Sigma-LAG fractions were not the most active (Fig. 4A). Within the classical-nucleation framework, these differences correspond to variation in effective contact angle and therefore in the ability of the molecular surface to support formation of an ice nucleus (Fig. 4D). The warmer INA of some pollen INMs is consistent with their presenting more favourable ice-nucleating surfaces than Sigma-LAG, although effective contact angle here represents various unresolved molecular and structural properties rather than a directly measured material constant.

The importance of molecular aggregation is also supported by the Yariv reagent experiments. Binding of β-glucosyl Yariv reagent to Sigma-LAG increased its median freezing temperature by 4.2 °C (Fig. 3C). Because Yariv reagent associates with sufficiently long β-1,3-galactan chains, this result indicates that changing the association state of Type II AG can enhance its INA. Conversely, the peak in activity among the size-fractionated Sigma-LAG samples shows that increasing molecular size or aggregation is not sufficient by itself: the organisation and surface properties of the resulting assemblies must also matter.

Type II AGs are highly branched and structurally variable, with differences in backbone length, side-chain composition and molecular association both within and among samples (*36, 42, 50*). We therefore propose that their INA emerges from this structural heterogeneity rather than from a single ice-nucleating structure present in every molecule. This is consistent also with our observation that a structurally homogeneous mycobacterial AG did not nucleate ice effectively (Fig. 3A). A solution can contain a very large number of AG molecules and assemblies, only a small proportion of which need to adopt structures capable of nucleating ice at the warmest observed temperatures (Figs. 4C and D). We refer to this as a “chance nucleator” model: increasing molecular size makes warm nucleation physically possible, while chance structural variation determines how frequently an assembly presents a sufficiently large and favourable ice-nucleating surface (Fig. 4C).

This model provides a possible explanation for both the widespread occurrence and striking variability of plant INA. Arabinogalactans are ancient, ubiquitous and structurally heterogeneous components of land plants, and their association with ice nucleation in pollen, fern spores and moss spores therefore suggests that this activity is an incidental property of these molecules rather than a specialised function that evolved independently in each group. In this respect, arabinogalactans resemble many other ice nucleators, including AgI and feldspars, whose ice-nucleating activity is obviously incidental rather than adaptive. Arabinogalactan differs, however, in lacking an obvious structural complementarity with ice. The heat-resistant activity commonly observed between approximately –14 and –30 °C in PWW and SWW may arise when rare AG molecules or assemblies become sufficiently large and suitably organised to nucleate ice. Variation in AG composition and association among samples would alter the frequency of these chance nucleators, producing substantial differences in INA even among related plant species. The warmer, heat-sensitive activity of some PWW and SWW samples may arise when proteins organise AG chains into larger or more favourable assemblies. This interpretation is consistent with the effects of molecular size, Yariv-induced association and heat treatment, but the structures responsible for the highest-temperature nucleation remain to be determined. It does not exclude subsequent selection for ice nucleation in particular lineages, but removes the need to invoke such selection to explain its broad phylogenetic distribution.

### Environmental implications of plant arabinogalactan ice nucleation

Fern spores can occur in the atmosphere over vegetated regions at abundances comparable to pollen grains (*51*). Their contribution to atmospheric ice nucleation should therefore be considered alongside that of pollen, although their atmospheric abundance and importance as INPs remain poorly constrained. Moss spores too are wind dispersed (*52*) although atmospheric abundance has not been assessed to our knowledge. Beyond propagules, AGs also occur widely in plant tissues and may enter the atmosphere through the fragmentation, dispersal and degradation of plant material. Related polysaccharides are produced by phylogenetically diverse algae (*53*), raising the possibility that AG-like molecules also contribute to marine organic aerosol.

More broadly, our results suggest that structurally variable, soluble macromolecules represent an important and underappreciated route to atmospheric ice nucleation. We identify Type II arabinogalactans as the likely molecular basis of the soluble INMs released by diverse plant materials. Their ubiquity and solubility favour broad environmental distribution, while variation in their molecular size and supramolecular organisation may explain the highly variable INA observed among plant-derived samples.

## Author contributions

Conceptualization: NLHK, TFW

Methodology: NLHK, WGTW, DB, MIG, GCS, TFW

Investigation: NLHK, EC, FB, AB, DES, MID, CIB, IL, CF, CSG, ANB, DB, MIG, GCS, TFW

Formal analysis: NLHK, GCS, TFW

Visualization: NLHK, GCS, TFW

Funding acquisition: DB, MIG, GCS, TFW

Project administration: TFW

Supervision: WGTW, MIG, GCS, TFW

Writing – original draft: NLHK, TFW

Writing – review & editing: NLHK, EC, FB, AB, DES, MID, CIB, IL, CF, CSG, ANB, JPK, WGTW, DB, MIG, GCS, TFW.

## Diversity, equity, ethics, and inclusion

This study did not involve human participants, human-derived data, or Indigenous knowledge. Plant materials were obtained from commercial suppliers or collected in accordance with applicable institutional and local requirements. The composition of the research team reflected the expertise required to conduct the study.

## Data, code, and materials availability

All data, simulation input files and code supporting the findings of this study will be deposited in the University of Leeds Research Data Repository (https://archive.researchdata.leeds.ac.uk/). The repository DOI will be added before publication.

## References

1. T. Koop, B. J. Murray, A physically constrained classical description of the homogeneous nucleation of ice in water. J. Chem. Phys. 145, 211915 (2016).

2. D. A. Knopf, P. A. Alpert, Atmospheric ice nucleation. Nat. Rev. Phys. 5, 203–217 (2023).

3. A. Korolev et al., Mixed-phase clouds: Progress and challenges. Meteorol. Monogr. 58, 5.1–5.50 (2017).

4. S. M. Burrows et al., Ice-nucleating particles that impact clouds and climate: Observational and modeling research needs. Rev. Geophys. 60, e2021RG000745 (2022).

5. R. J. Herbert et al., Gaps in our understanding of ice-nucleating particle sources exposed by global simulation of the UK Earth System Model. Atmos. Chem. Phys. 25, 291–325 (2025).

6. M. Z. Jacobson, D. G. Streets, Influence of future anthropogenic emissions on climate, natural emissions, and air quality. J. Geophys. Res. Atmos. 114, D08118 (2009).

7. S. Werchner et al., When do subpollen particles become relevant for ice nucleation processes in clouds? J. Geophys. Res. Atmos. 127, e2021JD036340 (2022).

8. K. Diehl, S. Matthias-Maser, R. Jaenicke, S. K. Mitra, The ice nucleating ability of pollen: Part II. Laboratory studies in immersion and contact freezing modes. Atmos. Res. 61, 125–133 (2002).

9. N. von Blohn, S. K. Mitra, K. Diehl, S. Borrmann, The ice nucleating ability of pollen: Part III. New laboratory studies in immersion and contact freezing modes including more pollen types. Atmos. Res. 78, 182–189 (2005).

10. B. G. Pummer, H. Bauer, J. Bernardi, S. Bleicher, H. Grothe, Suspendable macromolecules are responsible for ice nucleation activity of birch and conifer pollen. Atmos. Chem. Phys. 12, 2541–2550 (2012).

11. S. Augustin et al., Immersion freezing of birch pollen washing water. Atmos. Chem. Phys. 13, 10989–11003 (2013).

12. E. Gute, J. P. D. Abbatt, Oxidative processing lowers the ice nucleation activity of birch and alder pollen. Geophys. Res. Lett. 45, 1647–1653 (2018).

13. E. Gute, J. P. D. Abbatt, Ice nucleating behavior of different tree pollen in the immersion mode. Atmos. Environ. 231, 117488 (2020).

14. J. Burkart, J. Gratzl, T. M. Seifried, P. Bieber, H. Grothe, Isolation of subpollen particles (SPPs) of birch: SPPs are potential carriers of ice nucleating macromolecules. Biogeosciences 18, 5751–5765 (2021).

15. N. L. H. Kinney, C. A. Hepburn, M. I. Gibson, D. Ballesteros, T. F. Whale, High interspecific variability in ice nucleation activity suggests pollen ice nucleators are incidental. Biogeosciences 21, 3201–3214 (2024).

16. B. H. Matthews, A. N. Alsante, S. D. Brooks, Pollen emissions of subpollen particles and ice nucleating particles. ACS Earth Space Chem. 7, 1207–1218 (2023).

17. K. Dreischmeier, C. Budke, L. Wiehemeier, T. Kottke, T. Koop, Boreal pollen contain ice-nucleating as well as ice-binding “antifreeze” polysaccharides. Sci. Rep. 7, 41890 (2017).

18. J. Kretzschmar et al., From trees to rain: Enhancement of cloud glaciation and precipitation by pollen. Environ. Res. Lett. 19, 104052 (2024).

19. Y. Zhang et al., Effects of pollen on hydrometeors and precipitation in a convective system. J. Geophys. Res. Atmos. 129, e2023JD039891 (2024).

20. M. C. Wozniak, F. Solmon, A. L. Steiner, Pollen rupture and its impact on precipitation in clean continental conditions. Geophys. Res. Lett. 45, 7156–7164 (2018).

21. M. Prank, J. Tonttila, X. Shang, S. Romakkaniemi, T. Raatikainen, Can pollen affect precipitation? Atmos. Chem. Phys. 25, 183–197 (2025).

22. P. Wolber, G. Warren, Bacterial ice-nucleation proteins. Trends Biochem. Sci. 14, 179– 182 (1989).

23. J. Fröhlich-Nowoisky et al., Ice nucleation activity in the widespread soil fungus *Mortierella alpina*. Biogeosciences 12, 1057–1071 (2015).

24. H. J. Tong et al., A new electrodynamic balance (EDB) design for low-temperature studies: Application to immersion freezing of pollen extract bioaerosols. Atmos. Meas. Tech. 8, 1183–1195 (2015).

25. L. Felgitsch et al., Birch leaves and branches as a source of ice-nucleating macromolecules. Atmos. Chem. Phys. 18, 16063–16079 (2018).

26. B. J. Harris et al., Divergent evolutionary trajectories of bryophytes and tracheophytes from a complex common ancestor of land plants. *Nat*. Ecol. Evol. 6, 1634–1643 (2022).

27. P. K. Strother, C. Foster, A fossil record of land plant origins from charophyte algae. Science 373, 792–796 (2021).

28. S. Wallace, A. Fleming, C. H. Wellman, D. J. Beerling, Evolutionary development of the plant spore and pollen wall. AoB Plants 2011, plr027 (2011).

29. T. F. Whale, Quantification of the ice nucleation activity of ice-binding proteins using a microliter droplet freezing experiment. Methods Mol. Biol. 2730, 121–134 (2024).

30. B. G. Pummer et al., Ice nucleation by water-soluble macromolecules. Atmos. Chem. Phys. 15, 4077–4091 (2015).

31. M. Kačuráková, P. Capek, V. Sasinková, N. Wellner, A. Ebringerová, FT-IR study of plant cell wall model compounds: Pectic polysaccharides and hemicelluloses. Carbohydr. Polym. 43, 195–203 (2000).

32. M. I. Daily, M. D. Tarn, T. F. Whale, B. J. Murray, An evaluation of the heat test for the ice-nucleating ability of minerals and biological material. Atmos. Meas. Tech. 15, 2635–2665 (2022).

33. J. L. Morris et al., The timescale of early land plant evolution. Proc. Natl. Acad. Sci. U.S.A. 115, E2274–E2283 (2018).

34. P. C. J. Donoghue, C. J. Harrison, J. Paps, H. Schneider, The evolutionary emergence of land plants. Curr. Biol. 31, R1281–R1298 (2021).

35. X. Ma, C. Zhang, L. Yang, S. B. Hedges, B. Zhong, New insights on angiosperm crown age based on Bayesian node dating and skyline fossilized birth-death approaches. Nat. Commun. 16, 2265 (2025).

36. B. Classen, A. Baumann, J. Utermoehlen, Arabinogalactan-proteins in spore-producing land plants. Carbohydr. Polym. 210, 215–224 (2019).

37. T. M. Seifried et al., Fluorescence signal of proteins in birch pollen distorted within its native matrix: Identification of the fluorescence suppressor quercetin-3-*O*-sophoroside. Anal. Bioanal. Chem. 414, 7531–7542 (2022).

38. F. Wieland et al., Aggregation of ice-nucleating macromolecules from *Betula pendula* pollen determines ice nucleation efficiency. Biogeosciences 22, 103–115 (2025).

39. D. Bartels et al., Evolution of plant cell wall: Arabinogalactan-proteins from three moss genera show structural differences compared to seed plants. Carbohydr. Polym. 163, 227–235 (2017).

40. A. Clarke, R. Anderson, B. Stone, Form and function of arabinogalactans and arabinogalactan-proteins. Phytochemistry 18, 521–540 (1979).

41. A. Majewska-Sawka, E. A. Nothnagel, The multiple roles of arabinogalactan proteins in plant development. Plant Physiol. 122, 3–10 (2000).

42. A. M. Showalter, Arabinogalactan-proteins: Structure, expression and function. Cell. Mol. Life Sci. 58, 1399–1417 (2001).

43. Y. Ma, K. Johnson, Arabinogalactan proteins: Multifunctional glycoproteins of the plant cell wall. Cell Surf. 9, 100102 (2023).

44. K.-K. Mueller, L. Pfeifer, L. Wegner, K. Ehlers, B. Classen, New insights into bryophyte arabinogalactan-proteins from a hornwort and a moss model organism. Plant J. 123, e70312 (2025).

45. T. A. Torode et al., Branched pectic galactan in phloem-sieve-element cell walls: Implications for cell mechanics. Plant Physiol. 176, 1547–1558 (2018).

46. T. L. Lowary, Twenty years of mycobacterial glycans: Furanosides and beyond. Acc. Chem. Res. 49, 1379–1388 (2016).

47. K. Kitazawa et al., β-Galactosyl Yariv reagent binds to the β-1,3-galactan of arabinogalactan proteins. Plant Physiol. 161, 1117–1126 (2013).

48. T. F. Whale, T. Surawy-Stepney, S. L. Barr, Survival-analysis methods for droplet-freezing data: Kaplan–Meier confidence intervals and non-parametric curve comparison. Atmos. Meas. Tech. 19, 6699–6718 (2026).

49. N. H. Fletcher, Size effect in heterogeneous nucleation. J. Chem. Phys. 29, 572–576 (1958).

50. V. A. Babkin, N. A. Neverova, E. N. Medvedeva, T. E. Fedorova, A. A. Levchuk, Investigation of physicochemical properties of arabinogalactan of different larch species. Russ. J. Bioorg. Chem. 42, 707–711 (2016).

51. V. R. Després et al., Primary biological aerosol particles in the atmosphere: A review. Tellus B Chem. Phys. Meteorol. 64, 15598 (2012).

52. B. van Zanten, Experimental studies on trans-oceanic long-range dispersal of moss spores in the Southern Hemisphere. J. Hattori Bot. Lab. 44, 455–482 (1978).

53. C. Hervé et al., Arabinogalactan proteins have deep roots in eukaryotes: Identification of genes and epitopes in brown algae and their role in *Fucus serratus* embryo development. New Phytol. 209, 1428–1441 (2016).

54. T. F. Whale et al., A technique for quantifying heterogeneous ice nucleation in microlitre supercooled water droplets. Atmos. Meas. Tech. 8, 2437–2447 (2015).

55. C. I. Biggs et al., Mimicking the ice recrystallization activity of biological antifreezes: When is a new polymer “active”? Macromol. Biosci. 19, 1900082 (2019).

56. D. M. Murphy, T. Koop, Review of the vapour pressures of ice and supercooled water for atmospheric applications. Q. J. R. Meteorol. Soc. 131, 1539–1565 (2005).

57. D. Barahona, Analysis of the effect of water activity on ice formation using a new thermodynamic framework. Atmos. Chem. Phys. 14, 7665–7680 (2014).

58. E. M. Goellner, J. Utermoehlen, R. Kramer, B. Classen, Structure of arabinogalactan from *Larix laricina* and its reactivity with antibodies directed against type-II arabinogalactans. Carbohydr. Polym. 86, 1739–1744 (2011).

59. S.-J. Park et al., CHARMM-GUI Glycan Modeler for modeling and simulation of carbohydrates and glycoconjugates. Glycobiology 29, 320–331 (2019).

60. O. Guvench et al., CHARMM additive all-atom force field for carbohydrate derivatives and its utility in polysaccharide and carbohydrate–protein modeling. J. Chem. Theory Comput. 7, 3162–3180 (2011).

61. J. L. F. Abascal, E. Sanz, R. García Fernández, C. Vega, A potential model for the study of ices and amorphous water: TIP4P/Ice. J. Chem. Phys. 122, 234511 (2005).

62. C. M. Miles, P.-C. Hsu, A. M. Dixon, S. Khalid, G. C. Sosso, Lipid bilayers as potential ice nucleating agents. Phys. Chem. Chem. Phys. 24, 6476–6491 (2022).

63. R. Chandrasekaran, S. Janaswamy, Morphology of Western larch arabinogalactan. Carbohydr. Res. 337, 2211–2222 (2002).

64. A. Barducci, G. Bussi, M. Parrinello, Well-tempered metadynamics: A smoothly converging and tunable free-energy method. Phys. Rev. Lett. 100, 020603 (2008).

65. M. Bonomi et al., Promoting transparency and reproducibility in enhanced molecular simulations. Nat. Methods 16, 670–673 (2019).

66. M. Salvalaglio, P. Tiwary, M. Parrinello, Assessing the reliability of the dynamics reconstructed from metadynamics. J. Chem. Theory Comput. 10, 1420–1425 (2014).

67. D. van der Spoel et al., GROMACS: Fast, flexible, and free. J. Comput. Chem. 26, 1701–1718 (2005).

68. M. A. Cuendet, W. F. van Gunsteren, On the calculation of velocity-dependent properties in molecular dynamics simulations using the leapfrog integration algorithm. J. Chem. Phys. 127, 184102 (2007).

69. S. Miyamoto, P. A. Kollman, SETTLE: An analytical version of the SHAKE and RATTLE algorithm for rigid water models. J. Comput. Chem. 13, 952–962 (1992).

70. B. Hess, P-LINCS: A parallel linear constraint solver for molecular simulation. J. Chem. Theory Comput. 4, 116–122 (2008).

71. G. Bussi, D. Donadio, M. Parrinello, Canonical sampling through velocity rescaling. J. Chem. Phys. 126, 014101 (2007).

72. M. Bernetti, G. Bussi, Pressure control using stochastic cell rescaling. J. Chem. Phys. 153, 114107 (2020).

73. G. A. Tribello, F. Giberti, G. C. Sosso, M. Salvalaglio, M. Parrinello, Analyzing and driving cluster formation in atomistic simulations. J. Chem. Theory Comput. 13, 1317–1327 (2017).

74. P. J. Steinhardt, D. R. Nelson, M. Ronchetti, Bond-orientational order in liquids and glasses. Phys. Rev. B 28, 784–805 (1983).

75. P. Rodger, T. Forester, W. Smith, Simulations of the methane hydrate/methane gas interface near hydrate-forming conditions. Fluid Phase Equilib. 116, 326–332 (1996).

76. F. Bachtiger, T. R. Congdon, C. Stubbs, M. I. Gibson, G. C. Sosso, The atomistic details of the ice recrystallisation inhibition activity of PVA. Nat. Commun. 12, 1323 (2021).

77. R. J. Gowers et al., “MDAnalysis: A Python package for the rapid analysis of molecular dynamics simulations,” in Proceedings of the 15th Python in Science Conference, S. Benthall, S. Rostrup, Eds. (SciPy, Austin, TX, 2016), pp. 98–105.

78. C. P. Garnham, R. L. Campbell, P. L. Davies, Anchored clathrate waters bind antifreeze proteins to ice. Proc. Natl. Acad. Sci. U.S.A. 108, 7363–7367 (2011).

79. C. F. Weber, *Polytrichum commune* spores nucleate ice and associated microorganisms increase the temperature of ice nucleation activity onset. Aerobiologia 32, 353–361 (2016).

80. B. F. Moffett, Ice nucleation in mosses and liverworts. Lindbergia 38, 14–16 (2015).

81. J. Yariv, H. Lis, E. Katchalski, Precipitation of arabic acid and some seed polysaccharides by glycosylphenylazo dyes. Biochem. J. 105, 1C–2C (1967).

82. M. Jermyn, Y. Yeow, E. Woods, A class of lectins present in the tissues of seed plants. Aust. J. Plant Physiol. 2, 501–531 (1975).

83. A. Clarke, P. Gleeson, M. Jermyn, R. Knox, Characterization and localization of β-lectins in lower and higher plants. Aust. J. Plant Physiol. 5, 707–722 (1978).

84. POWO, Plants of the World Online (Royal Botanic Gardens, Kew, 2026); https://powo.science.kew.org/ (accessed 5 November 2026).

85. Tropicos.org, Missouri Botanical Garden, St. Louis, MO; https://www.tropicos.org/ (accessed 5 November 2026).

86. J.C. Brinda, J.J. Atwood, Eds., The Bryophyte Nomenclator; https://www.bryonames.org/ (accessed 5 November 2026).

