## Supplementary materials for "Arabinogalactans provide the molecular basis of ice nucleation by plant pollen and spores"

Nina L. H. Kinney *et al.*

**The PDF file includes:**

Materials and Methods
Supplementary Text
Figs. S1 to S18
Tables S1 to S2
References

### Materials and Methods

#### Experimental Methods

##### Fern Spore Collection

Fern spores were collected first by identifying fertile fronds with filled sporangia (spore sacks found on the underside of the fronds). The fertile fronds were removed from the plant and placed sporangia-side down on paper to dry out, allowing dehiscence of the sporangia. Once the spores were released onto the paper, they were sieved, transferred to sealed containers, and refrigerated. The time of spore collections and the locations and species of the parent ferns are detailed in Table S1.

##### Moss Spore Collection

Mature moss capsules were harvested from sporulating specimens into sample bags. The spores were extracted from capsules, using clean tweezers, directly into glass vials and stored at 4 °C until use. Specimen collection details are included in Table S2.

##### Spore Solution Preparation

Each spore suspension was prepared by immersing a known mass of spores in Milli-Q® water in a clean glass vial (approximately 0.04 g spores in 2 mL water for 2 wt% suspensions). The suspension was shaken by hand to disperse the spores in the water and refrigerated for approximately 24 hours before filtration. To prepare the spore washing water (SWW) solutions, the suspension was filtered using a 0.2 μm syringe filter into a clean glass vial, removing the spores and insoluble fragments, as shown in Fig. S1.

##### Pollen and Polysaccharide Sample Details

European hornbeam (*Carpinus betulus* L.), silver birch (*Betula pendula* Roth), elder (*Sambucus nigra* L.) and ponderosa pine (*Pinus ponderosa* Douglas ex C. Lawson) pollen samples were obtained from Pharmallerga. Mycobacterial arabinogalactan was obtained through BEI Resources, NIAID, NIH: Mycobacterium tuberculosis, Strain H37Rv, Purified Arabinogalactan, NR-14852. Arabinogalactan from larch wood obtained from abcr GmbH is labelled as 'abcr-LAG' and arabinogalactan from larch wood obtained from Sigma-Aldrich labelled as 'sigma-LAG' throughout. β-glucosyl Yariv reagent and gum arabic were obtained from Biosupplies Australia.

##### Ice Nucleation Measurements and Analysis

Most Ice nucleation measurements were conducted using a Peltier-driven microlitre droplet freezing experiment (μ-DFE), described in detail by Whale et al. (*29*). Briefly, a Hampton Research siliconised glass slide, holding an array of approximately 50 × 1 μL sample solution droplets, was placed on the temperature-controlled stage and the temperature lowered at a rate of 1 °C/min. A camera was used to monitor the freezing temperatures of droplets. Two independent freezing runs were conducted for each sample solution. The Sigma-AG data shown as black squares in Fig. 4A were obtained using the μL-NIPI instrument described by Whale et al. (*54*). This instrument follows the same measurement principle and experimental procedure as the Peltier-driven μ-DFE described above, but uses an Asymptote EF600 Stirling cryocooler rather than a Peltier element to control the stage temperature.

Figures indicating the ice-nucleating activity (INA) of various solutions display the fraction of frozen droplets as a function of temperature, $f_{ice}(T)$, given by

$$f_{ice}(T) = \frac{n(T)}{N}, \qquad \text{S1}$$

where $n(T)$ is the number of frozen droplets and $N$ is the total number of droplets.

For soluble ice nucleators, the cumulative number of nucleation sites per unit dry mass (e.g. number of nucleation sites per gram spores) $n_m(T)$, can be calculated from the fraction of droplets frozen, $f_{ice}(T)$, and the mass of ice nucleator in or contributing to each droplet of the sample solution, $M$, by the following equation:

$$n_m(T) = \frac{-\ln(1 - f_{ice}(T))}{M}. \qquad \text{S2}$$

Pointwise confidence intervals were calculated using the Kaplan–Meier method, implemented in Python (*48*). The Kaplan–Meier estimate of the fraction of droplets remaining unfrozen is

$$\hat{S}(T) = \prod_{T_j \geq T} \left[\frac{N_U(T_j) - N_F(T_j)}{N_U(T_j)}\right], \qquad \text{S3}$$

where $N_U(T_j)$is the number of unfrozen droplets immediately before temperature $T_j$, and $N_F(T_j)$ is the number freezing at that temperature. The fraction frozen is then $f(T) = 1 - \hat{S}(T)$. The variance of the log–log-transformed Kaplan–Meier estimate was calculated as

$$\hat{\sigma}^2(T) = \frac{1 - \hat{S}(T)}{N_0 \hat{S}(T)[\ln \hat{S}(T)]^2}. \qquad \text{S4}$$

The confidence limits for $\hat{S}(T)$were then calculated as

$$\left[\hat{S}(T)^{\exp\left(+z_{\alpha/2}\hat{\sigma}(T)\right)}, \hat{S}(T)^{\exp\left(-z_{\alpha/2}\hat{\sigma}(T)\right)}\right], \qquad \text{S5}$$

where $z_{\alpha/2} = 1.96$ for a 95% confidence interval. Confidence limits for $f(T)$ were obtained by subtracting the upper and lower limits for $\hat{S}(T)$ from 1, and were propagated to $n_m(T)$ using the equation above. These are pointwise confidence intervals rather than simultaneous confidence bands across the complete temperature range.

<u>Comparison of freezing curves using the Integrated Squared Difference test</u>

Freezing curves were compared using a permutation-based Integrated Squared Difference (ISD) test recently described by Whale et al. (*48*). The test statistic is the integral of the squared separation between two fraction-frozen curves over temperature. Significance was determined by randomly reallocating observations between groups while preserving sample sizes and recalculating the statistic for each permutation. Unlike tests based solely on shifts in central tendency, the ISD test is sensitive to differences in both the position and shape of freezing curves. In this analysis, 10,000 random permutations were performed. Consequently, the minimum attainable p-value was approximately $1 \times 10^{-4}$. Where no permuted statistic equaled or exceeded the observed statistic, the result was therefore reported as $p < 1 \times 10^{-4}$.

Fourier Transform Infrared Spectroscopy

For the commercially supplied arabinogalactan (AG) from larch wood (samples obtained from abcr GmbH labelled 'abcr-LAG' and from Sigma-Aldrich labelled 'Sigma-LAG'), pectin from Sigma-Aldrich and gum arabic from Biosupplies, FT-IR measurements were conducted using unprocessed dry samples. For collections of pollen, spore and larch bark samples, 20 wt% suspensions were prepared and refrigerated for 24 hours before 0.2 µm filtration. The filtrates were freeze dried and the FT-IR spectra recorded for each of the freeze-dried residues. FT-IR measurements were conducted using an Agilent Technologies Cary 630 FT-IR Spectrometer.

Heat Treatment

Heat tests were performed according to the wet heating method described by Daily et al. (*32*), considered sufficient to denature proteinaceous ice nucleators. Ice nucleation measurements of untreated PWWs, fern and moss SWWs were conducted first, using the µ-DFE. 1 mL of each of the solutions was then transferred into a screw top vial which was sealed tightly, and the vial placed in a boiling water bath on a heat block for exactly 30 minutes. The volume of the water bath was maintained above the liquid level of the solutions by adding small amounts of room temperature water at intervals, so the pollen and fern spore solutions were fully immersed in the boiling water throughout. After allowing the vials to cool to room temperature, ice nucleation measurements were conducted for the heat-treated solutions.

Aqueous Size Exclusion Chromatography

Size Exclusion Chromatography (SEC) analysis of AG, pectin and gum arabic was performed on an Agilent Technologies 1260 instrument equipped with differential refractive index (DRI), light scattering (LS) and viscometry (VS) detectors. The columns used were 1x PL aquagel-OH guard, 1x PL aquagel-OH 30, 1x PL aquagel-OH 40 and 1x PL aquagel OH 60. The eluant was 0.1 M $NaNO_3$ in water/methanol 80/20. Samples were dissolved in the eluant at 3 mg/mL and filtered using 0.2 µm Millipore® hydrophilic filters.

Nuclear Magnetic Resonance Spectroscopy (NMR)

Solutions of abcr-LAG and Sigma-LAG were prepared at a concentration of 20 mg/mL AG in $D_2O$. NMR spectra were recorded on a Bruker Avance III HD 400 MHz, with $D_2O$ as the solvent. Chemical shifts of protons are reported as δ in parts per million (ppm) relative to the solvent residual peak of $D_2O$ at 4.79 ppm.

Yariv Reagent Tests

The impact of β-glucosyl Yariv reagent on nucleation temperature was measured for six samples: purified water, gum arabic, abcr-LAG, Sigma-LAG, silver birch (Betula pendula) pollen, and elder (*Sambucus nigra*) pollen.

In each case, 100 µL of 0.3 M NaCl solution was combined with 100 µL of 0.02 wt% sample solution e.g. 0.02 wt% abcr-LAG solution. The combined 200 µL 0.01 wt% sample 0.15 M NaCl solution was then sub-divided into two 100 µL aliquots. The first was tested directly on the μ-DFE, without the addition of Yariv reagent. The second was combined with 50 μg of Biosupplies β-glucosyl Yariv reagent, shaken, and left for 30 minutes at room temperature before testing on the μ-DFE. The fraction of droplets frozen for each of these solutions was compared to evaluate the impact of β-glucosyl Yariv reagent binding on ice nucleation ability.

Ice Recrystallisation Inhibition Assay and Analysis

Ice recrystallisation inhibition properties of solutions were evaluated using the 'splat' assay as described by Biggs et al. (*55*). Approximately 10 μL of the sample in phosphate-buffered saline (PBS) solution was ejected from a syringe held 1.40 m above a glass coverslip atop a thin aluminium plate, cooled to approximately –78 °C on dry ice. On impact with the glass coverslip, the solution droplet spread out, forming an approximately 10 μm thin ice layer. The glass coverslip was immediately transferred onto a Linkam cryostage and held at –8 °C to anneal for 30 minutes. Images were captured, using a Canon EOS 500D DSLR camera set above an Olympus CX41 microscope fitted with a UIS-2 20x/0.45/∞/0-2/FN22 lens and Olympus crossed polarisers, immediately after placing the sample in the Linkam stage, to ensure a poly-crystalline ice monolayer had formed, and then again after 30 minutes.

The number of ice crystals in the field of view photographed at the 30-minute mark was counted in ImageJ and divided by the area of the field of view. The mean of three independent measurements for each sample concentration was calculated to give the mean grain size (MGS) which was compared with that of the control PBS solution to give MGS as a percentage relative to PBS.

Ice Shaping Measurements

Ice shaping measurements were conducted for three solutions separately: 200 mg/mL AG in 30 wt% sucrose solution, 20 μM Type III antifreeze protein (positive control) in 30 wt% sucrose solution, and 30 wt% sucrose solution (negative control) using an Otago Osmometers nanolitre osmometer. For each solution, several ~20 nL droplets were suspended in Type B Cargille immersion oil on a 6-well cooling plate via a microcapillary system. The sample droplets were then cooled rapidly until all were frozen, around –30 °C. The temperature was then increased at a rate of 0.01 °C/min until a single ice crystal remained in a single droplet. The temperature was gradually lowered again until crystal growth was clearly visible. The observations were captured with a Canon EOS 1200D digital SLR camera and an Olympus CX41 microscope fitted with a UIS-2 20x/0.45/∞/0-2/FN22 lens.

Microarray polymer profiling

Aqueous extracts of plant materials and polysaccharide standards were printed as microarrays onto nitrocellulose membrane with a pore size of 0.45 μm (Whatman) using an Arrayjet Sprint (Arrayjet, Roslin, Scotland). Each sample was spotted at three concentrations (starting at neat aqueous extracts or 1 mg/mL in the case of polysaccharide standards and followed by two fivefold dilutions) and as three technical replicates. After printing, the microarrays were probed with monoclonal antibodies or a carbohydrate binding module (CBM27) which were detected using appropriate anti-mouse, anti-rat or (in the case of CBM27) anti-his secondary antibodies conjugated to alkaline phosphatase. The microarrays were developed using an NBT/BCIP substrate containing nitroblue tetrazolium (NBT) and 5-bromo-4-chloro-3-indolylphosphate p-toluidine salt (BCIP) in alkaline phosphatase buffer (100 mM NaCl, 5 mM MgCl2, 100 mM diethanol-amine, pH 9.5). The arrays were scanned using a desktop scanner (CanoScan 8800F, Canon) at 2400 dpi and spot intensities determined using microarray analysis software (Array-Pro Analyzer 6.3 (Media Cybernetics). The highest value in the data sets was assigned a value of 100 and all other signals normalised accordingly.

**Calculation of size- and contact angle-dependent heterogeneous ice nucleation temperatures**

Fig. 4B shows the temperature at which heterogeneous ice nucleation on a spherical particle reaches an observable rate of $J_{\mathrm{het}} = 1\ \mathrm{cm^{-2}s^{-1}}$. The rate was calculated using classical nucleation theory:

$$J_{\mathrm{het}} = \frac{k_{\mathrm{B}}T}{h} n_1 \exp\left(-\frac{\Delta G_{\mathrm{diff}}}{k_{\mathrm{B}}T}\right) \exp\left(-\frac{f_{\mathrm{het}}\Delta G_{\mathrm{crit}}}{k_{\mathrm{B}}T}\right), \qquad \mathrm{S6}$$

where $n_1 = 10^{15}\ \mathrm{cm^{-2}}$.

The temperature-dependent diffusion activation free energy was calculated using the power-law expression of Koop and Murray (*1*),

$$\Delta G_{\mathrm{diff}}(T) = k_{\mathrm{B}}T/2 + \gamma k_{\mathrm{B}}T^2/(T - T_{\mathrm{s}}), \qquad \mathrm{S7}$$

with the best-fit parameters $T_{\mathrm{s}} = 215.45$ K and $\gamma = 1.9188$. This expression gives $\Delta G_{\mathrm{diff}}$ in joules per molecule and accounts for the rapid reduction in the translational mobility of water molecules with decreasing temperature. The free-energy barrier that must be overcome to form a stable, critically sized ice nucleus in supercooled water was calculated as

$$\Delta G_{\mathrm{crit}}(T) = \frac{16\pi v_{\mathrm{i}}^2(T)\sigma_{\mathrm{iw}}^3(T)}{3[\Delta\mu(T)]^2}. \qquad \mathrm{S8}$$

The thermodynamic driving force was calculated from the Murphy and Koop (*56*) vapour-pressure parameterisations for supercooled liquid water and ice:

$$\Delta\mu(T) = k_{\mathrm{B}}T \ln\left(\frac{p_{\mathrm{w}}(T)}{p_{\mathrm{i}}(T)}\right). \qquad \mathrm{S9}$$

To determine the interfacial tension between water and ice, Barahona's temperature and water activity dependent parameterisation was used (*57*).

$$\sigma_{\mathrm{iw}} = 0.00211 - 0.0513a_{\mathrm{w}} + 3.04 \times 10^{-4}T, \qquad \mathrm{S10}$$

where $T$ is in kelvin and $\sigma_{\mathrm{iw}}$ is in $\mathrm{J\ m^{-2}}$.

The effects of particle size and contact angle were represented using the Fletcher (*49*) expression for nucleation on a finite spherical particle:

$$f_{\mathrm{het}} = \frac{1}{2}\left[1 + \left(\frac{1-mX}{\phi}\right)^3 + X^3\left\{2 - 3\left(\frac{X-m}{\phi}\right) + \left(\frac{X-m}{\phi}\right)^3\right\} + 3mX^2\left\{\frac{X-m}{\phi} - 1\right\}\right], \qquad \mathrm{S11}$$

where $X = r_{\mathrm{NP}}/r^*$, $m = \cos\theta$, and $\phi = (1 + X^2 - 2Xm)^{1/2}$. Here, $r_{\mathrm{NP}}$ is the radius of the spherical nucleating particle, $r^*$is the temperature-dependent critical ice-cluster radius, and $\theta$ is the contact angle. Particle mass was converted to $r_{\mathrm{NP}}$assuming spherical particles with a density of $2000\ \mathrm{kg\ m^{-3}}$ (appropriate for carbohydrates) and particle mass is displayed in kilodaltons. Because $r^*$varies with temperature, $X$and $f_{\mathrm{het}}$were recalculated at each temperature.

For each combination of particle mass and contact angle, the nucleation rate was evaluated between 225.0 and 273.14 K. Brent's root-finding method was used to identify the warmest temperature at which $J_{\mathrm{het}} = 1\ \mathrm{cm^{-2}s^{-1}}$. Each contour therefore joins combinations of particle mass and contact angle predicted to reach the reference nucleation rate at the same temperature. The selected rate provides a common basis for comparison and is not intended to represent a universal experimental detection threshold.

#### Computer Simulations

A molecular model for LAG

We built a molecular model of larch arabinogalactan (LAG) based on the structural information reported in (*58*). The polysaccharide comprises 75 sugar residues: 63 galactose (Gal) residues, including a β-(1→3)-D-galactan backbone with β-(1→6)-linked galactosyl side chains; 9 α-Larabinofuranose (Ara) residues, present as Ara-(1→6)-Gal and Ara-(1→3)-Ara-(1→6)-Gal motifs; and 3 β-D-glucuronic acid (GlcA) residues attached through (1→6) linkages to Gal. The resulting sequence is illustrated in Fig. S15.

It is important to note that this model represents only a small segment of the much larger, complex network of (cross-linked and/or aggregate) LAG actually found in nature. The aim of utilising this model is to investigate water and ice in close proximity to these sugars, not capture the impact on ice nucleation of structural features on a broader, longer scale.

We utilised CHARMM-GUI Glycan Modeler (*59*) to generate a topology for this structure according to the CHARMM36 force field (*60*). In terms of the water model, we opted for TIP4P/Ice (*61*), based on a number of previous works which demonstrated the suitability of utilising the CHARMM36 force field in conjunction with the TIP4P/Ice water model (see e.g., Ref. (*62*)).

The LAG model obtained via CHARMM-GUI is shaped into a helical structure. This is consistent with previous molecular models of Western LAG(*63*). However, we wanted to explore the conformational space of our relatively short helix. To this end, we adopted metadynamics (*64*) simulations, utilising as collective variable the radius of gyration, $R_g$, of the polysaccharide backbone. The PLUMED software(*65*), version 2.9.2, has been used to drive these enhanced sampling simulations. The relevant parameters are: Gaussian height (0.6 kJ/mol), Gaussian width (0.05), deposition stride (2 ps), and bias factor (500). The details of the molecular dynamics (MD) simulations underpinning this technique are reported in the next section. The uncertainty associated with the resulting free energy profile has been computed via the reweighing technique described in Ref (*66*).

The result is reported in Fig. S16, and indicates that, possibly due to the relatively short length of the polysaccharide chain and the drive to maximise its intra-molecular (HB) bonds, our LAG model seems to favour a configuration where the helix, whilst still apparent, is characterised by a much shorter pitch. Said pitch was estimated to be 5.19 ± 0.23 Å, computed as half the axis-projected separation between equivalent LAG ring centres separated by 11 backbone residues, corresponding to two turns of the helix. The resulting configuration is approximately spherical in shape. Note that these metadynamics simulations have been performed to explore the conformational space of the LAG model, not to compute an accurate free energy profile. In fact, the large bias factor utilised makes these well-tempered metadynamics simulations almost equivalent to standard metadynamics.

<u>MD simulations</u>

MD simulations were performed via the GROMACS MD package (*67*), version 2024.4-CUDA-12.4.0. The choice of the relevant force fields is discussed in the previous section. The initial dimension of the (cubic) simulation box is 14 nm in the x, y and z directions. The system comprises the 75-sugar LAG model described in the previous section, as well as ~90,162 water molecules and 3 $Na^+$ ions, to balance the negative charge due to the three GlcA [(1→6)-$\beta$-D-glucoronic acid] residues (which are typically deprotonated in physiological and/or atmospheric conditions). The equations of motion were propagated via a leapfrog integrator(*68*) with a timestep of 2 fs. The atomic positions were saved every 40 ps. Periodic boundary conditions were applied in all Cartesian directions. The cutoff for the van der Waals and electrostatic interactions was set to 12 and 10 Å, respectively: a switching function was used to bring the van der Waals interaction to zero at 12 Å. The geometry of the water molecules was constrained using the SETTLE algorithm (*69*)while the P-LINCS algorithm (*70*)was used to constrain the bonded C/O–H terms in the LAG to their equilibrium value. The MD simulations were performed within the NPT ensemble, utilising the Bussi-Donadio-Parrinello thermostat (*71*) and the Bernetti-Bussi barostat (isotropic conditions, ambient pressure) (*72*), with coupling constants of 1 and 2 ps, respectively.

To investigate the emergence of pre-critical ice nuclei, we first equilibrated the system at 300 K for 50 ns. The temperature was then lowered to 230 K over 50 ns, which corresponds to a linear cooling ramp of 1.4 K/ns. The statistics for ice nuclei presented here are averaged over 2,500 configurations, sampled every 40 ps throughout a 100 ns production run at 230 K and ambient pressure following the cooling ramp. We performed five statistically independent replicas of these simulations by initialising the starting velocities at 300 K from the corresponding Maxwell–Boltzmann distribution using five different random seeds. In combination with the stochastic thermostat and barostat, this procedure ensures that the replicas sample independent realisations of the same thermodynamic protocol.

<u>Identifying ice nuclei</u>

To identify the pre-critical ice nuclei, we employed the PLUMED package (*65*) – specifically version 2.11.0-dev, which enables the analysis of a large number of water/ice molecules. The details of the approach underpinning the methodology we used can be found in Ref. (*73*) to identify "ice-like" clusters by combining (i) hydrogen-bond topology, (ii) Steinhardt orientational order parameters ($Q_6$)(*74*), and (iii) short-range proximity, via leveraging adjacency/contact matrices and performing a clustering of "icy" water molecules via depth-first search (*73*). We begin by constructing a hydrogen bond (HB) contact matrix for the water molecules within our system, with the donor–hydrogen–acceptor geometry controlled by Gaussian switching functions on distances (oxygen-oxygen distance of 3.2 Å) and angles (oxygen-hydrogen-oxygen angle between 150 and 180 degrees). This returns an *NxN* adjacency matrix *hb_mat* whose (*i,j*) element is the (smoothed) number/weight of HBs between molecules *i* and $j$. To count the HBs per molecule, we right-multiply hb_mat by a vector of ones and select molecules with exactly four HBs (which yields the vector hfilt). Next, we compute the LOCAL_Q6 parameter (*75*) *l6(i)* for each water molecule in the system. This parameter is constructed from the Steinhardt order parameter $Q_6$ as:

$$q_{6m}(i) = \frac{\sum_{j\neq i}\sigma(|r_{ij}|)Y_{6m}(\hat{r}_{ij})}{\sum_{j\neq i}\sigma(|r_{ij}|)},$$

$$\hat{q}_{6m}(i) = \frac{q_{6m}(i)}{\left(\sum_{m=-6}^{6}|q_{6m}(i)|^2\right)^{1/2}},$$

$$\ell_6(i) = \frac{\sum_{j\neq i}\sigma(|r_{ij}|)\sum_{m=-6}^{6}\hat{q}_{6m}(i)\hat{q}_{6m}^{*}(j)}{\sum_{j\neq i}\sigma(|r_{ij}|)} \qquad \text{S12}$$

where $q_{6m}(i)$ and $q_{6m}(j)$ are the sixth order Steinhardt vectors calculated for atom $i$ and atom $j$ respectively and the asterisk denotes complex conjugation. The function $\sigma(\boldsymbol{rij})$ is a switching function that acts on the distance between atoms $i$ and $j$. The vector of $l_6(i)$ values that is obtained from this calculation is transformed using a switching function, resulting in a vector, cf , whose elements are all between 0 and 1. This vector can be used to identify the molecules that are part of the ice nucleus as the switching function ensures that element $i$ is only one if $l_6(i)$ >0.44, which is the threshold above which a water molecule is considered to be "icy") to obtain a smooth binary mask that upweights locally crystalline environments. We also built a conventional distance-based contact matrix cm, which encodes whether two molecules are within the solid-like first-neighbour shell. Note that all the oxygen-oxygen distance cutoff was chosen to be 3.24 Å, which is where the first minimum in the oxygen-oxygen pair correlation function for TIP4P/Ice water at 250 K is located (*76*). Next, we combined the information about the hydrogen bond network and the local order: we construct two outer-product masks: lq6_op = cf ⊗ cf and hb_f = hfilt ⊗ hfilt. We then compute the matrix, mat, as the following Hadamard product of these three matrices: mat = lq6_op X hb_f X cm. The *i,j* element of this matrix is thus only non-zero if (a) molecule $j$ is within the first coordination sphere of molecule $i$, (b) molecules $i$ and $j$ both participate in four hydrogen bonds and (c) molecules $i$ and $j$ are both in a part of the system where there is a high degree of local order. Finally, identify connected components of mat via a depth-first search algorithm (*73*) and write the indices of the largest cluster(s) to file. Thus, for each frame along our MD trajectories we can reliably identify (and sort by size) the pre-critical ice nucleus clusters.

We identified the 100 largest ice nuclei within each of the 2500 configurations of the system we considered. For each of these nuclei, we computed the minimum distance $d_{\min}$ as the minimum distance between any of the oxygen atoms within a given ice nucleus and any of the atoms within the LAG. Due to finite extent of the water phase surrounding the LAG model, one should expect $d_{\min} \rightarrow 0$ as the size of the ice nuclei increases (as there is only so much water available for a given ice nucleus to occur within the system). To quantify the extent of this bias, we computed the probability density function of $d_{\min}$ for a random distribution of "synthetic" ice nuclei of the same sizes of those we actually observed. Specifically, for each ice nucleus of a given size we randomly selected an oxygen within in our system constructed a synthetic analogue of the same size (i.e., the same number of molecules). For each of these synthetic nuclei we also computed $d_{\min}$ as well as the resulting probability density function.

Hydrogen bond analysis

In terms of possible hydrogen bonds (HB) between water and LAG, we have different possibilities: the water oxygens can act as HB donors, with the LAG oxygens (either -OH, C-O-C or $COO^-$ moieties) acting as acceptors. Or the LAG oxygens (-OH moiety) can act as donors with the water oxygen atoms acting as acceptors. We utilised MDAnalysis (*77*), version 2.7.0 to identify HB between the ice nuclei and the LAG. The relevant criteria are: (i.) the distance between acceptor and donor is less than or equal to 3.5 Å, and (ii.) the angle between donor-hydrogen-acceptor is greater than or equal to 150 degrees.

Water structuring at the interface with LAG

In order to assess whether the presence of the LAG significantly induces structural order within the interfacial water phase (or whether indeed is significantly disrupts the water hydrogen bond network), we computed the order parameter $F_4$, which quantifies the degree of tetrahedral hydrogen-bond organization by measuring the torsional alignment of adjacent water molecules. It is defined as $F_4 = \langle \cos(3\varphi) \rangle_\Omega$ , where $\langle \cdot \rangle_\Omega$ indicates an ensemble average across the first solvation shell Ω, and $\varphi$ denotes the H-O---O-H torsional angle (computed by choosing, on each molecule, the hydrogen furthest from the other oxygen). This approach was originally introduced in the context of identifying “clathrate-like” water(*75*). Clathrate hydrates are crystalline phases where water forms hydrogen-bonded cages around “guest” molecules; “clathrate-like”, (and, when surface-templated, “anchored clathrate”) water denotes such cage motifs, which have been proposed to aid ice nucleation in biological systems, particularly near ice nucleating proteins (*78*).

#### Supplementary Text

Ice Nucleation by Moss Spores

Bryophytes (mosses, liverworts and hornworts) are more distantly related to the pollen-producing seed plants than ferns, having diverged from vascular plants during the Cambrian period, 515–494 million years ago (*26*). About a decade ago, Weber (*79*) discovered that spores of the moss *Polytrichum commune* could nucleate ice at temperatures as warm as −15 °C. We collected spores from three moss species for ice nucleation and physicochemical characterization measurements to determine whether the ice nucleators associated with mosses (*79, 80*) are similar in nature to pollen and fern INMs.

Figure S4 shows the results of ice nucleation measurements for the moss spore samples. We found no substantial difference in freezing temperatures of unfiltered suspensions and 0.2 µm filtrates (SWWs) showing that the ice nucleating species is soluble and separates from the spores in water, as for fern spores and plant pollen. FT-IR absorbances for freeze-dried *Polytrichum formosum* SWW exhibited the same dominant polysaccharide absorbance peaks observed in fern spore and pollen extracts (Fig. 1(C)). Heat treatment of *Ceratodon purpureus* SWW selectively reduced the activity of the relatively rare ice-nucleating structures active above –15 °C, whereas the more abundant ice nucleators active below −15 °C were unaffected. This is a similar response to that found for *Sambucus nigra* PWW (Fig. 1 (B)). This behaviour suggests that the ice-nucleating structures active above approximately −15 °C depend upon a heat-sensitive proteinaceous constituent, whereas the structures responsible for nucleation below −15 °C are consistent with polysaccharide INMs.

Yariv Reactivity

The precipitation of AGs from aqueous plant extracts by Yariv phenylglycosides [1,3,5-tri(p-glycosyloxyphenylazo) 2,4,6 trihydroxybenzene] has been used historically to detect plant AGs in diverse taxa (*81-83*). It has since been established that Yariv reagents bind selectively to the β-1,3-galactan main chains characteristic of Type II AGs; with a β-1,3-galactan chain length of five residues necessary for aggregation and seven residues necessary for precipitation with Yariv reagent (*47*).

Results of ice nucleation measurements of pollen and LAG samples before and after treatment with β-glucosyl Yariv reagent are presented in Fig. S8. Fig. S8A shows that β-glucosyl Yariv reagent itself does not facilitate ice nucleation since there is no significant difference between ice nucleation temperatures in droplets of NaCl solution and droplets of β-glucosyl Yariv reagent in NaCl solution ($p = 0.388$).

A significant increase in INA with addition of β-glucosyl Yariv reagent is recorded for silver birch (*Betula pendula,* $p < 1 \times 10^{-4}$) PWW and Sigma-Aldrich LAG (Sigma-LAG, $p < 1 \times 10^{-4}$) solutions, shown in Fig. S8 (C) and (D) respectively. Our interpretation of these results, based on the findings of Kitazawa et al. (*47*) is that β-1,3-galactan chains between five and seven residues in length are present in *Betula pendula* and Sigma-LAG INMs. Aggregation of these INMs by β-glucosyl reagent binding to the β-1,3-galactan backbone increases the efficiency of ice nucleation. These results strongly support the role of Type II AGs in the INA of plant-derived materials.

There were no significant changes in INA with Yariv treatment of the high-activity elder (*Sambucus nigra*, $p = 0.067$) PWW and abcr LAG (abcr-LAG, $p = 0.4381$) solutions. Considering the specificity of Yariv binding to the β-1,3-galactan AG backbone and the established hypervariable nature of plant AG structures, differing responses in Yariv reactivity for the various solutions tested are unsurprising. The absence of an ice nucleation active aggregation response does not indicate that Type II AGs are absent from these extracts. Indeed, the AGP-containing gum arabic (Fig. S8(B)) sample, supplied as a positive control for binding (precipitation) with Yariv reagent, undergoes a small but significant ($p < 1 \times 10^{-4}$) decrease in INA with Yariv treatment.

Microarray polymer profiling

Multiple glycan epitopes were detected in aqueous extracts from pollen samples using monoclonal antibodies including arabinan (mAb LM6) homogalacturonan (mAb LM19), xyloglucan (mAbs LM15, LM25 and CCRC-M1), extensin (mAb LM1) and arabinogalactan protein (AGP, mAb JIM13). All mAbs showed expected specificities towards the polysaccharide standards. Regarding the pollen extracts, it is worth noting that apart from homogalacturonan, the polysaccharides bearing the detected epitopes are complex, branched polymers and in the case of extensin and AGP, heavily glycosylated. Glycans were not detected in extracts from bark or spores. It is possible that glycans were simply absent from these samples. However, the epitopes recognised by the mAbs used do not represent the full repertoire of structures that may be present and some glycans may have been undetected. It is also possible that low levels of glycans in bark and spores were below the detection limit of the technique.

FT-IR Analysis of Soluble Plant Extracts

FT-IR spectra were measured for freeze-dried pollen washing waters (PWWs) and spore washing waters (SWWs) and compared with spectra of ground *Larix decidua* bark, commercial AGs, gum arabic and pectin (Fig. 1C). The absorption spectra of the soluble pollen and spore extracts were consistent with those of the commercial AG samples.

Small variations are apparent in the fingerprint region, particularly between 1100 and 900 $cm^{-1}$, where ring vibrations and stretching vibrations of C–OH and C–O–C bonds overlap. Kacurakova et al. (*31*) identified a similar absorbance doublet in Type II AGs, attributing the dominant peak at approximately 1070 $cm^{-1}$ to galactopyranose (Galp) units and the lower-wavenumber peak at approximately 1040-1000 $cm^{-1}$ to arabinofuranose (Araf) residues in the side chains. The relative proportion of these units may therefore underlie the subtle differences observed between pollen and spore extracts. Development of a distinct lower-wavenumber component has previously been associated with reduced INA in pollen-derived extracts (*13*) and the same correlation is observed here: the least active spore and pollen samples (*Osmunda claytoniana* and *Pinus ponderosa*) and the smaller, less IN-active size fractions of *Carpinus betulus* PWW (Fig. S14) are associated with a lower-wavenumber shoulder near 1000 $cm^{-1}$. These observations are consistent with higher INA being associated with AG-like structures containing a greater relative abundance of galactopyranose backbone units relative to arabinofuranose side chains.

##### Characterisation of Commercial Arabinogalactans

Aqueous size exclusion chromatography (SEC) and NMR measurements were conducted to investigate differences between the commercial AG samples, abcr-LAG and Sigma-LAG. The SEC chromatograms (Fig. S9) show that the principal components of both materials have similar molecular weight distributions. However, light scattering measurements for abcr-LAG revealed an additional peak between 10 and 13 min retention time that was absent from Sigma-LAG, consistent with a small population of higher molecular weight species in the abcr-LAG sample, although the contribution of supramolecular organisation to this peak cannot be ruled out.

These observations are supported by the NMR measurements shown in Figs. S10-13. The 2D COSY spectrum of abcr-LAG contained a distinct cross peak at 4.0, 5.1 ppm that was absent from the Sigma-LAG spectrum, indicating structural differences between the two samples. Despite these differences, the FT-IR spectra of abcr-LAG and Sigma-LAG were effectively indistinguishable (Fig. 1C). Such variability is not unexpected given the structural diversity reported for AGs from different sources (*50*) and that FT-IR is sensitive to functional groups rather than connectivity. The SEC and NMR measurements suggest structural and compositional differences between the two commercial AG samples that are not resolved in FT-IR analysis and may account for the differences observed in their INA.

##### Ice Recrystallisation Inhibition and Ice Shaping

Ice recrystallisation inhibition (IRI) measurements for AG, pectin and *Betula pendula* pollen PWW are shown in Fig. S6. *Betula pendula* PWW inhibited ice recrystallisation to some extent, whereas both AG samples and pectin exhibited little to no IRI activity. The concentration reported for *B. pendula* PWW is based on the mass of pollen grains in the starting suspension and therefore likely underestimates the activity of the ice-binding molecules responsible for the observed response.

Dynamic ice-shaping measurements showed that both AG samples produced a flower-shaped ice crystal morphology (Fig. S7), similar to that previously reported for *B. pendula* pollen extracts (*17*). No shaping was observed for the sucrose control, whereas the Type III AFP control produced the distinct morphology associated with binding to the primary prism and pyramidal planes of ice. Together, these measurements demonstrate that the commercial AG samples bind ice and modify crystal growth but do not substantially inhibit ice recrystallisation. This lack of IRI activity is consistent with preferential binding to the basal plane of ice, whereas inhibition of ice recrystallisation is generally associated with binding to the more rapidly growing prism and pyramidal planes. The IRI activity observed in *B. pendula* PWW therefore suggests the presence of additional ice-binding molecules with different ice-plane binding characteristics.

Molecular Dynamics Simulations

To gain atomistic insight into the ice-nucleating ability of LAG, we performed molecular-dynamics simulations of a model LAG molecule in water. The computational details are discussed in the Materials and Methods section. The model was based on the structure reported in Ref. (*47*) and contained 75 sugar residues: 63 galactose residues [BGAL; (1→3)-β-D-galactose], nine arabinose residues [ALAR; α-L-arabinofuranose] and three glucuronic acid residues [BGLC; (1→6)-β-D-glucuronic acid] (Fig. S15). The LAG configuration obtained from metadynamics was placed in a cubic simulation box approximately 14 nm on each side to minimise finite-size effects. The system was then cooled to investigate the formation and spatial distribution of pre-critical ice-like clusters. Five statistically independent replicas were simulated using the same cooling protocol.

Across the five replicas, there was no strong or reproducible tendency for pre-critical ice-like clusters to form preferentially at the LAG–water interface (Fig. S17). A modest interfacial enrichment relative to a synthetic reference distribution was observed in only two replicas, and in both cases comparable or greater enrichment occurred elsewhere in the simulation box. The simulations therefore provide no consistent evidence that an isolated LAG molecule strongly enhances the local formation of pre-critical ice-like structures. The larger simulation volume necessarily makes complete sampling more difficult, but the absence of a consistent effect among the replicas suggests that any interfacial enhancement is small under the conditions examined.

Hydrogen-bond analysis showed that the simulated LAG molecule could interact directly with ice-like clusters, as expected for a hydrophilic polysaccharide containing numerous hydroxyl groups. However, these interactions did not result in preferential formation of ice-like clusters at the LAG surface. BGLC residues were disproportionately involved in hydrogen bonding between LAG and the ice-like clusters, despite only three such residues being present in the model structure (Fig. S15). This may reflect the presence of a deprotonated carboxylate group within these residues, which can readily participate in hydrogen-bonding interactions.

The simulated LAG strand adopted a helical configuration with a pitch $5.19 \pm 0.23$ Å. Although this indicates a degree of repeating supramolecular structure, we found no evidence that the helical arrangement promoted the formation of pre-critical ice-like clusters. The pitch is not characteristic of the nearest-neighbour O–O distances in ice, although it is close to the diameter of a hexagonal ring in the basal plane. The LAG conformation was obtained from metadynamics and should be regarded as a plausible model configuration rather than a definitive determination of the actual structure. The possible significance of the helical arrangement should therefore not be overinterpreted. It also remains possible that repeating structure becomes relevant only for clusters approaching the critical nucleus size, which could not be tested over the molecular sizes and timescales accessible to these simulations.

Several measures of water structure around LAG were also examined. These analyses were generally inconclusive. The most robust measure, the $F_4$ order parameter, showed only a marginal increase at the LAG–water interface, providing little evidence for substantial ordering of interfacial water by an isolated LAG molecule.

Overall, the simulations do not support a mechanism in which the surface chemistry or local hydration structure of an individual LAG molecule is sufficient to explain its experimentally observed ice-nucleating activity. Instead, the simulations are consistent with the experimental evidence that INA emerges at larger molecular sizes and depends on supramolecular association and organisation. Such assemblies are substantially larger and more structurally diverse than the isolated molecule represented here. The simulations therefore do not exclude the possibility that particular LAG assemblies present extended or favourably organised surfaces capable of nucleating ice. Rather, they support the proposed “chance nucleator” model, in which only a small proportion of sufficiently large and suitably organised structures possess strong ice-nucleating surfaces.

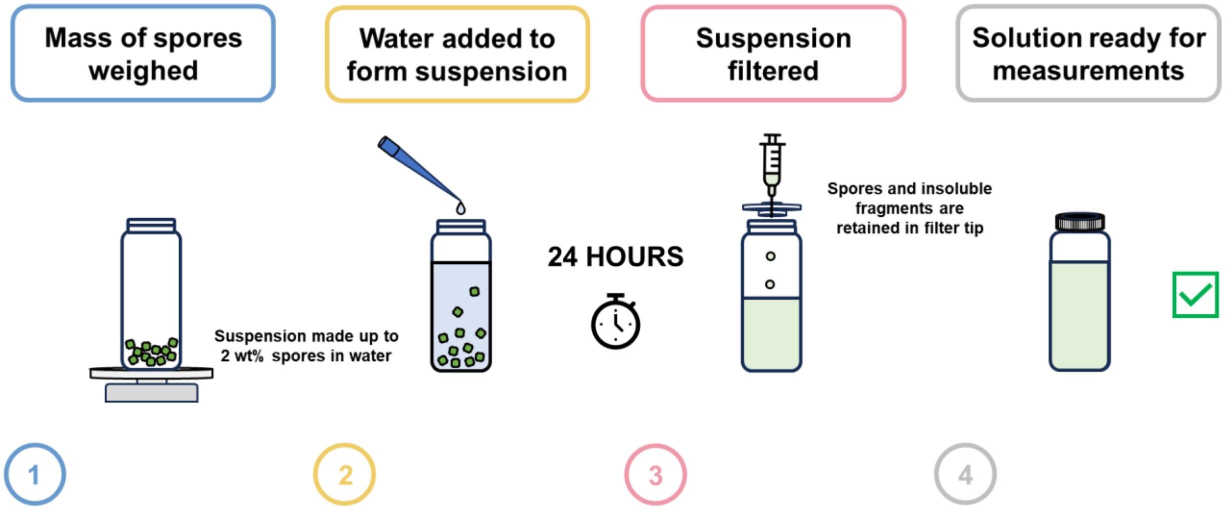


**Fig. S1.**
Schematic showing the preparation method for spore washing water (SWW) solutions, used for ice nucleation measurements.

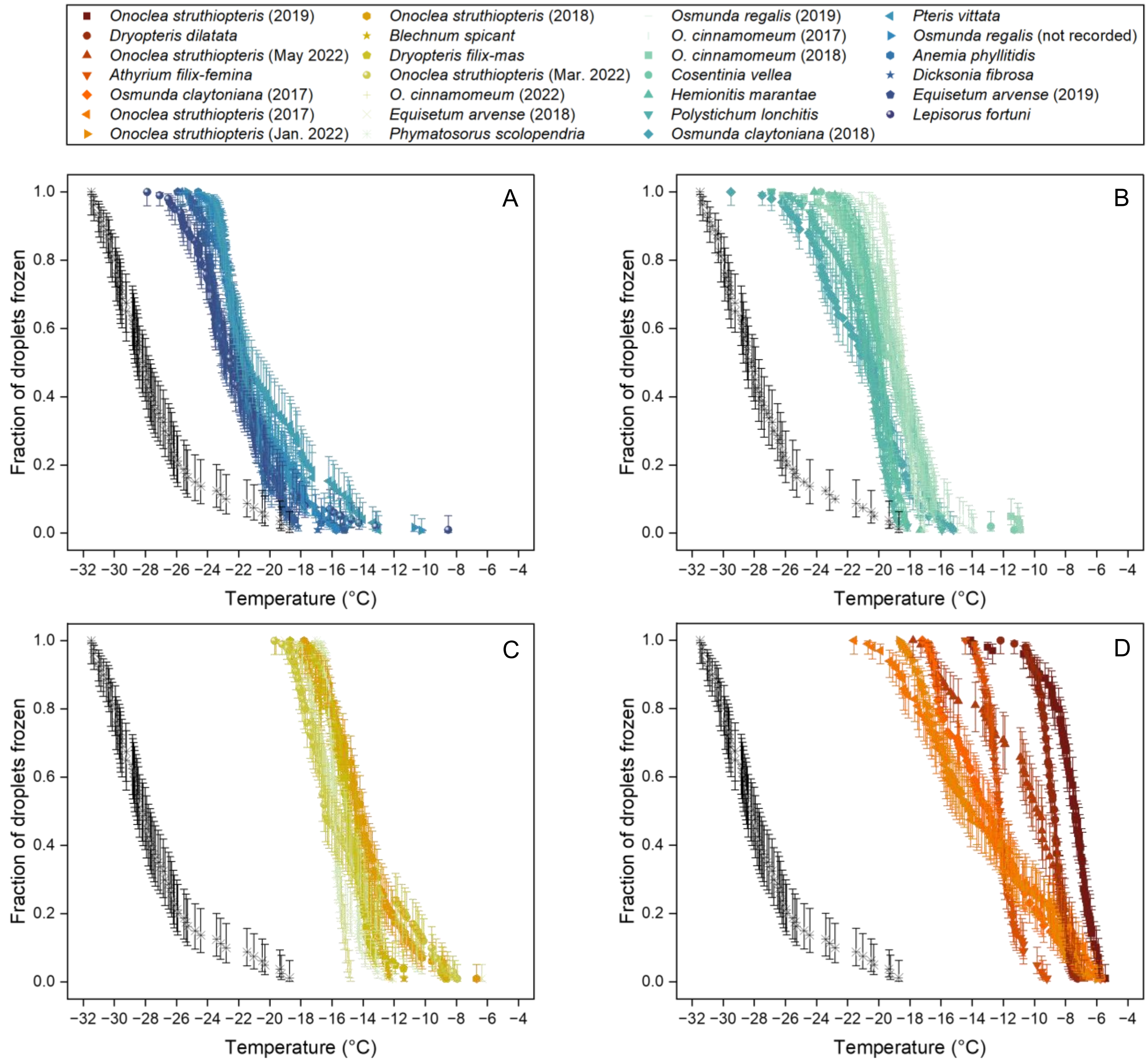


**Fig. S2.**
Fraction of 1 µL droplets of filtered fern spore solution frozen against temperature. Species name and collection date identify the spore samples tested. For clarity, the results are grouped into four panels: (**A**) (**B**) (**C**) and (**D**) based on ice nucleation activity, from low to high. Milli-Q® water is included in each as a background.

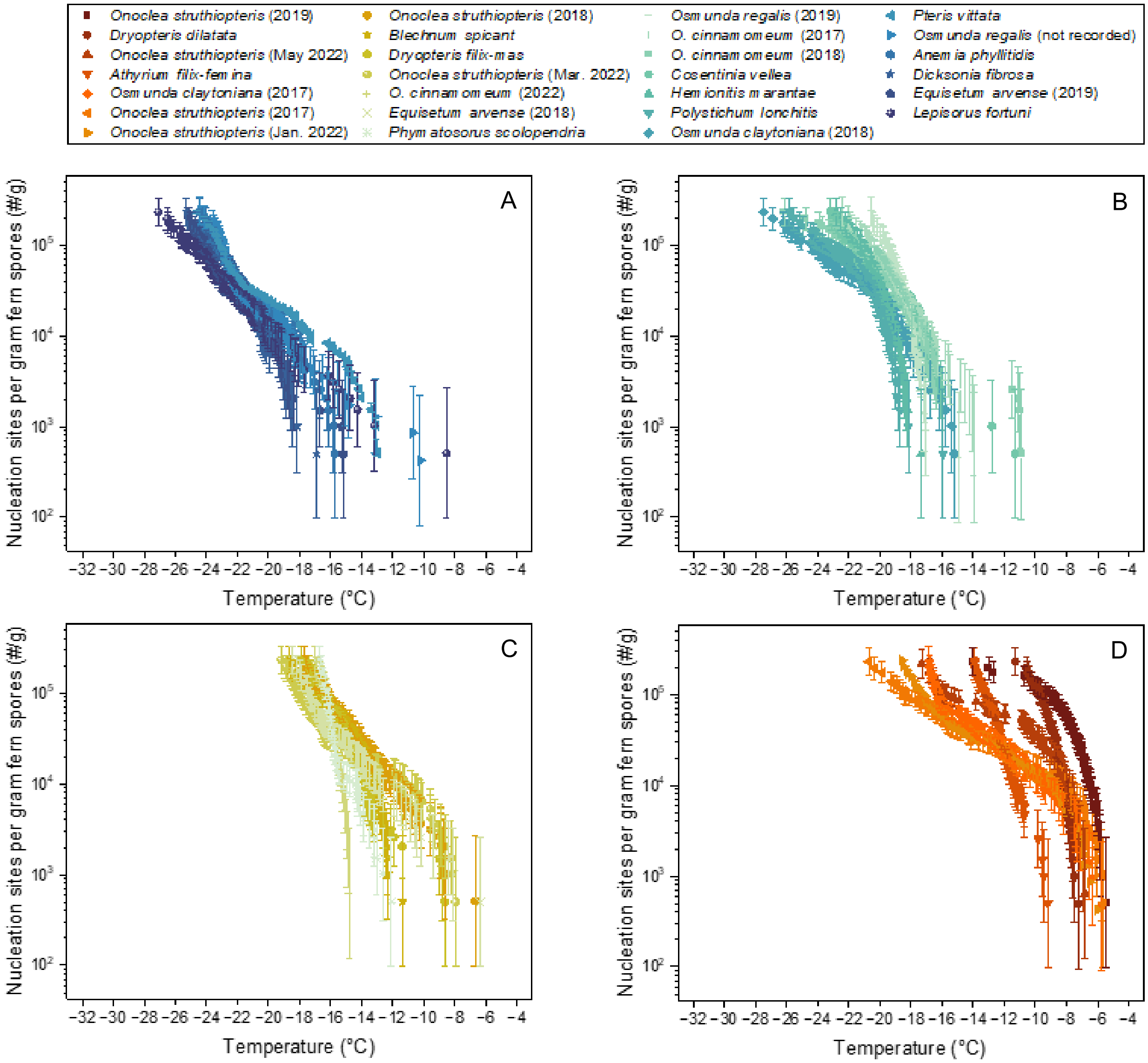


**Fig. S3.**
Number of nucleation sites per gram spores (g$^{-1}$) plotted against temperature for 1 μL droplets of filtered fern spore washing water solutions. Species name and collection date identify the spore samples tested. The results are grouped into four: (**A**) (**B**) (**C**) and (**D**) based on ice-nucleating activity, from low to high.

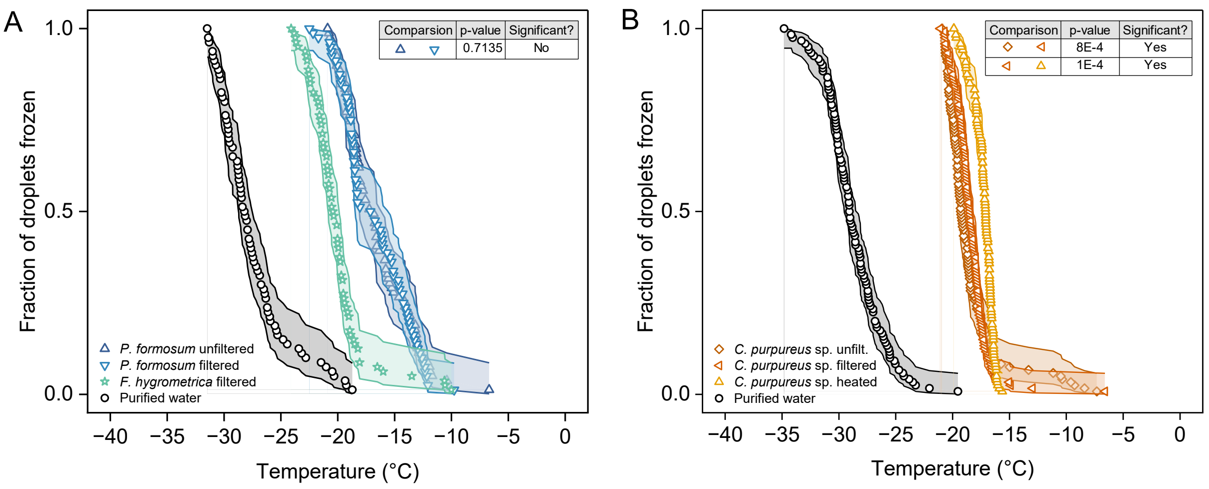


**Fig. S4.**

Fraction of 1 µL aqueous moss spore sample droplets frozen against temperature. The results are grouped by concentration of spores in the unfiltered starting suspensions: (**A**) 2 wt% *Polytrichum formosum* and *Funaria hygrometrica*, (**B**) 0.2 wt% *Ceratodon purpureus*, with the purified water instrument background included for each. 'Filtered' solutions (equivalent to spore washing water, SWW) were prepared by passing 'unfiltered' spore suspensions through 0.2 µm pore size filters. 'Heated' samples were wet heat-treated at 95 °C for 30 minutes, considered sufficient to inactivate proteinaceous ice nucleators (*32*). The significance of relevant comparisons, evaluated using the Integrated Squared Difference test (*48*) is tabulated.

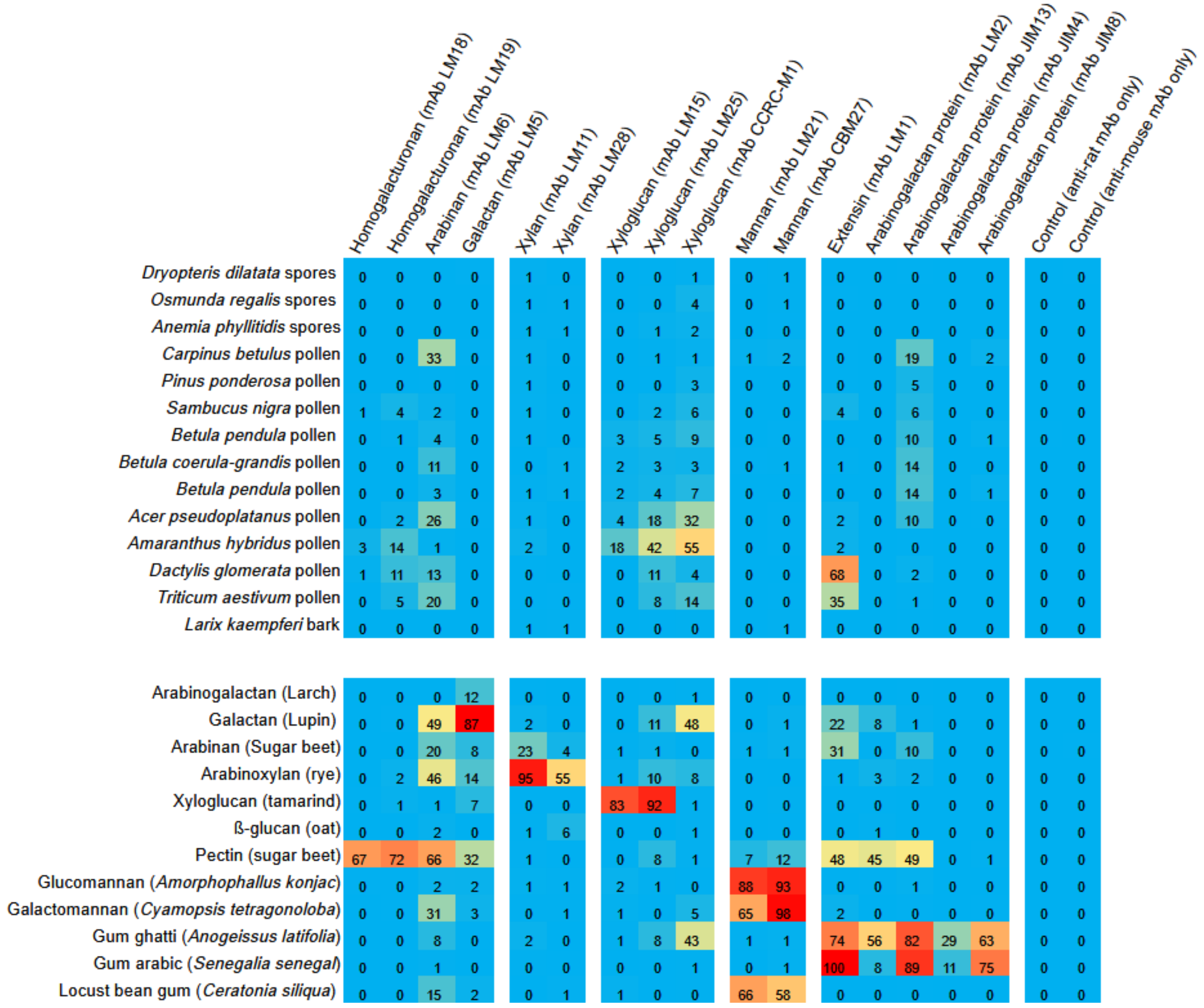


**Fig. S5.**

Heatmap showing the relative abundance of glycan structures (epitopes) detected in aqueous extracts of plant materials (spores, pollen and bark) and polysaccharide standards. Samples were printed onto nitrocellulose membrane as high-density microarrays which were then probed with monoclonal antibodies (mAbs, listed top) with specificity for glycan structures (epitopes). Values are mean spot signals. The highest value in the data sets was assigned a value of 100 and all other signals normalised accordingly.

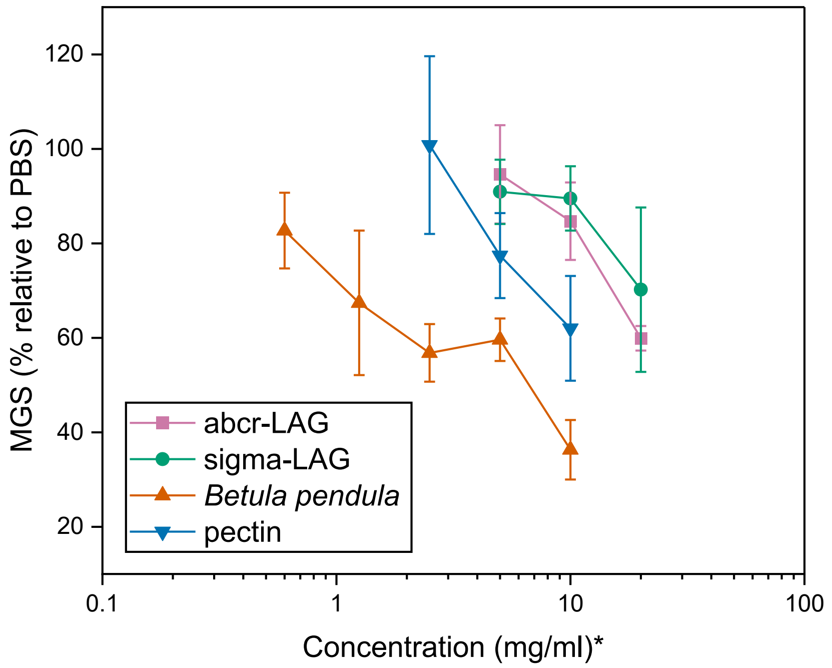


**Fig. S6.**
Comparison of ice recrystallisation activity, presented as mean grain size (MGS) as a percentage relative to that of phosphate-buffered saline (PBS), of abcr-LAG in PBS, Sigma-LAG in PBS, filtered *Betula pendula* pollen washing in PBS solution and pectin in PBS. *Note that the concentration in mg/mL used for the pollen washing solution is based on the mass of pollen in the starting suspension as the concentration of ice-binding molecules cannot be directly measured.

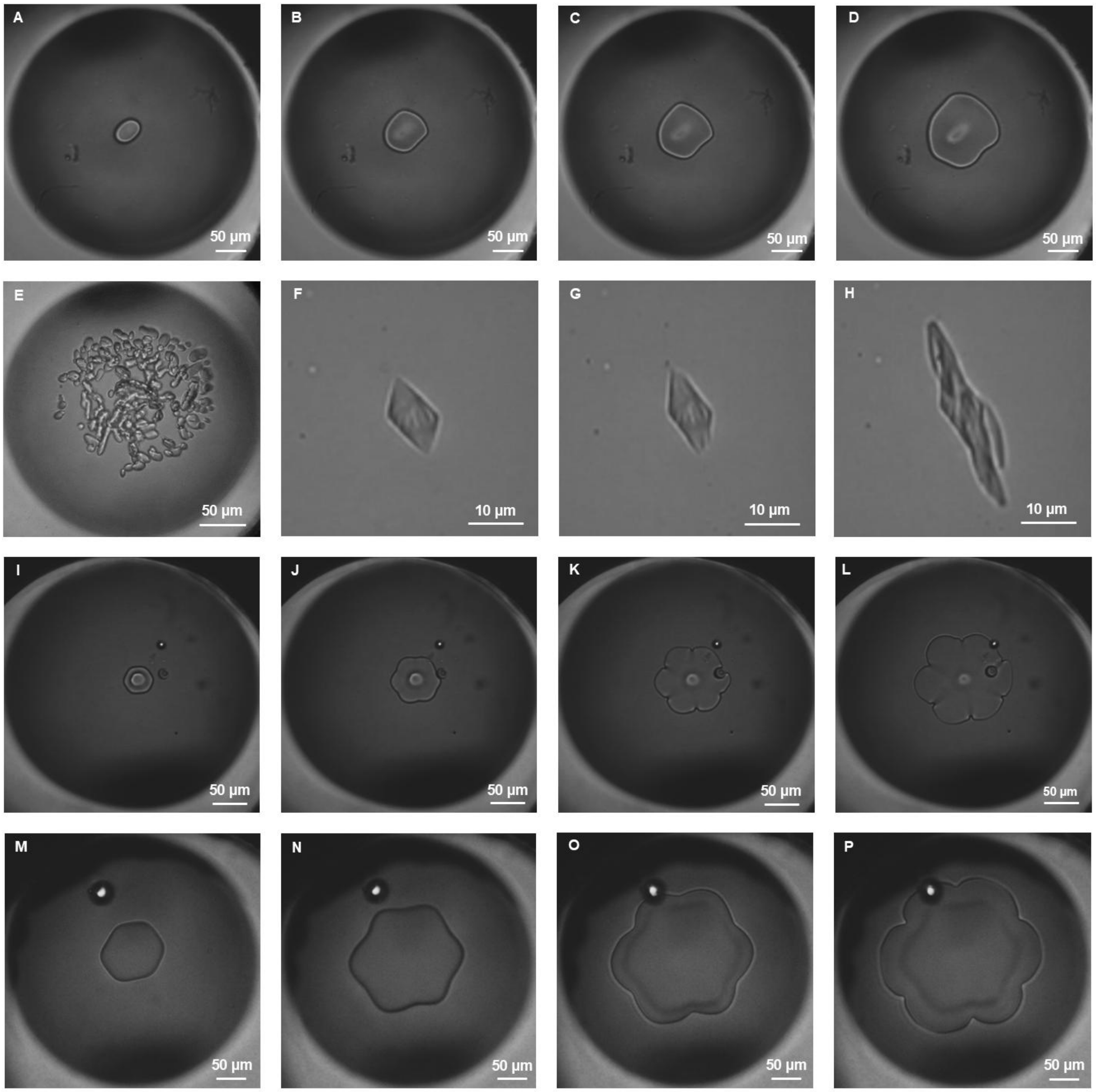


**Fig. S7.**

Results of ice shaping measurements: (**A-D**) show ice crystal growth in 30 wt% sucrose solution (negative control), (**E-H**) show ice crystal growth in solution of Type III antifreeze protein (primary prism plane and pyramidal plane ice binder) in 30 wt% sucrose, (**I-L**) show ice crystal growth in solution of Sigma-LAG in 30 wt% sucrose and (**M-P**) show ice crystal growth in solution of abcr-LAG in 30 wt% sucrose.

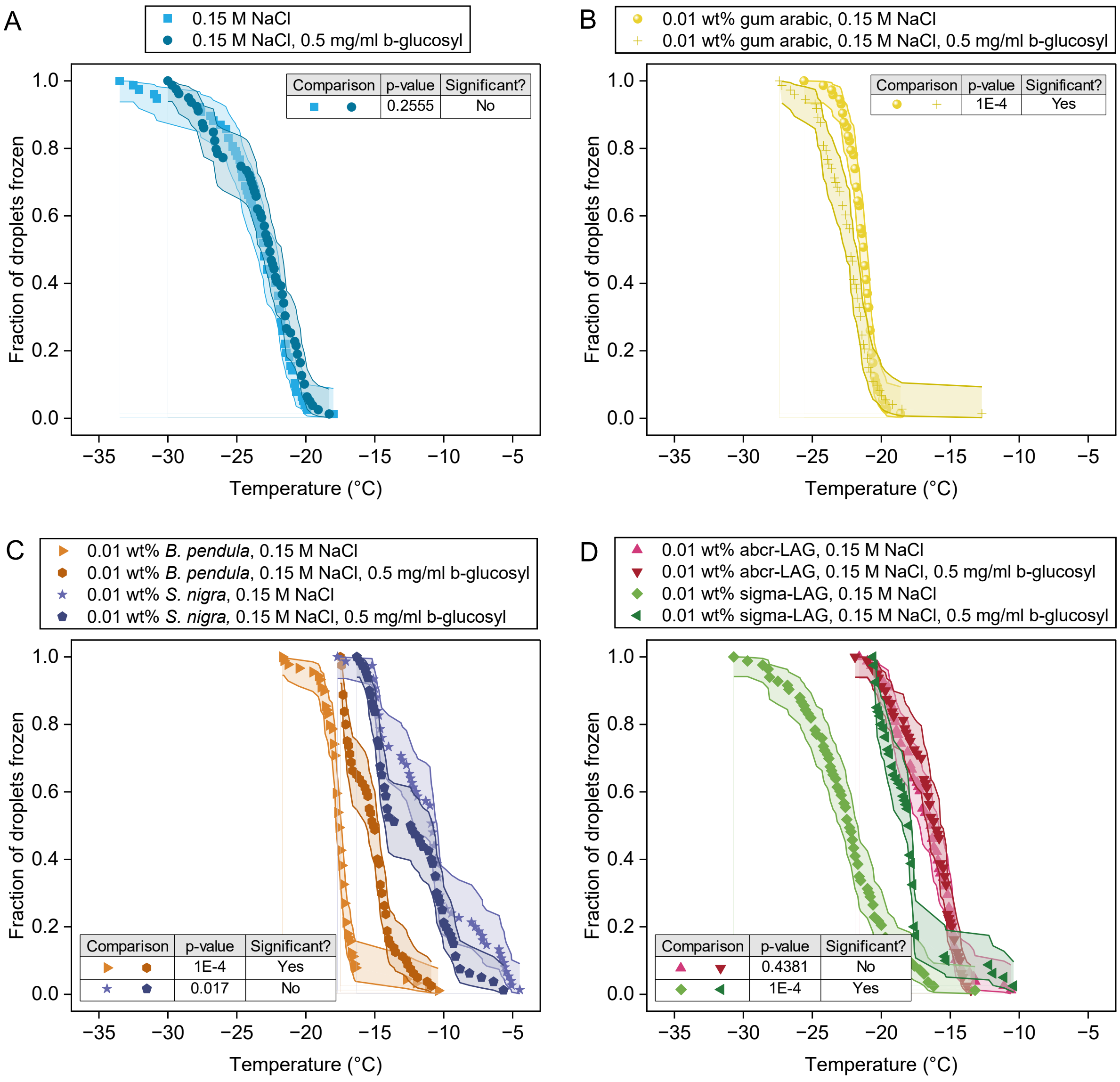


**Fig. S8.**

Fraction of 1 μL droplets frozen against temperature for various samples in 0.15 M NaCl solution, before and after the addition of β-glucosyl Yariv reagent. The samples tested were (**A**) a negative control solution of 0.15 M NaCl only, (**B**) 0.01 wt% Biosuppplies gum arabic, (**C**) filtered 0.01 wt% pollen washing water (PWW) solutions of silver birch (*Betula pendula*) and elder (*Sambucus nigra*) pollen obtained from Pharmallerga, (**D**) 0.01 wt% solutions of Sigma-Aldrich larch wood arabinogalactan (Sigma-LAG) and of abcr larch arabinogalactan (abcr-LAG).

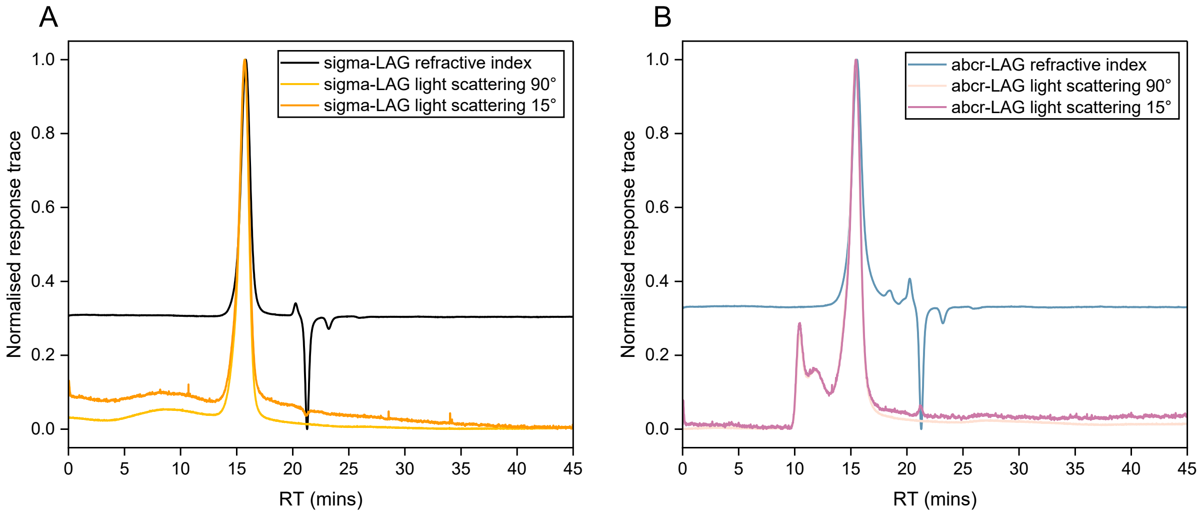


**Fig. S9.**

Results of aqueous size exclusion chromatography (SEC) measurements, comparing size distribution of larch wood arabinogalactan samples purchased from (**A**) Sigma-Aldrich (Sigma-LAG) and (**B**) from abcr GmbH (abcr-LAG). The columns used were 1x PL aquagel-OH guard, 1x PL aquagel-OH 30, 1x PL aquagel-OH 40 and 1x PL aquagel OH 60. The eluant was 80:20 0.1 M $NaNO_3$ (Aq):MeOH.

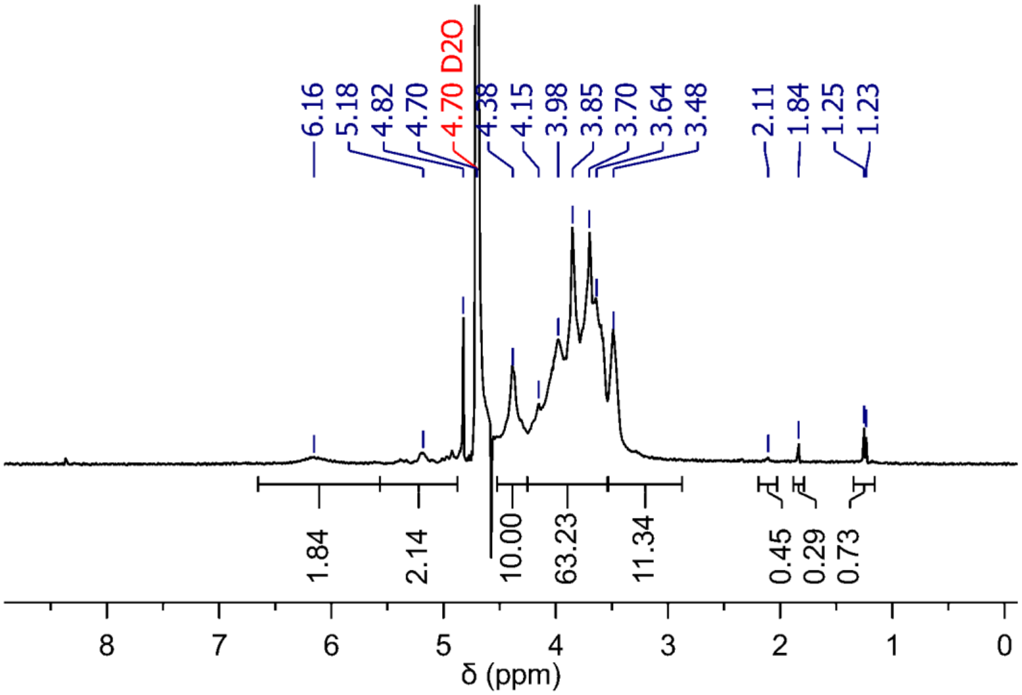


**Fig. S10.**
[1]H NMR spectrum ($D_2O$, 400 MHz) for larch wood arabinogalactan obtained from abcr GmbH (abcr-LAG).

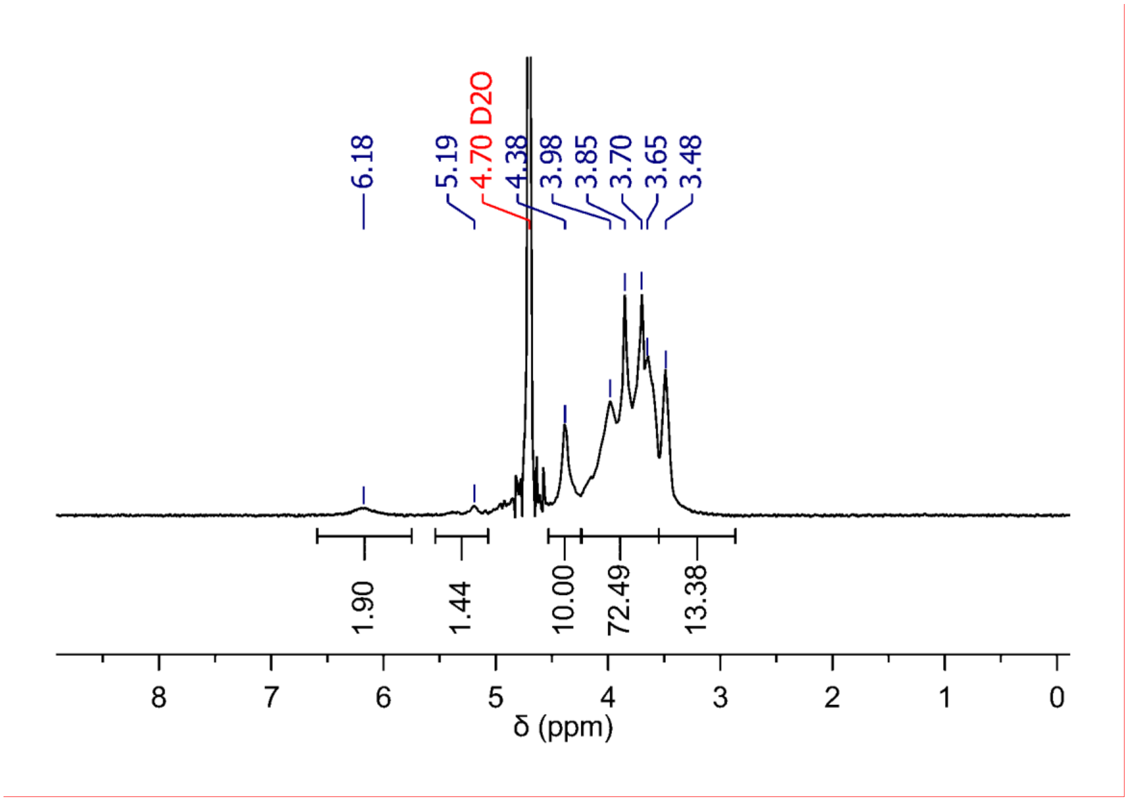


**Fig. S11.**

$^1$H NMR spectrum ($D_2O$, 400 MHz) for larch wood arabinogalactan obtained from Sigma-Aldrich (Sigma-LAG).

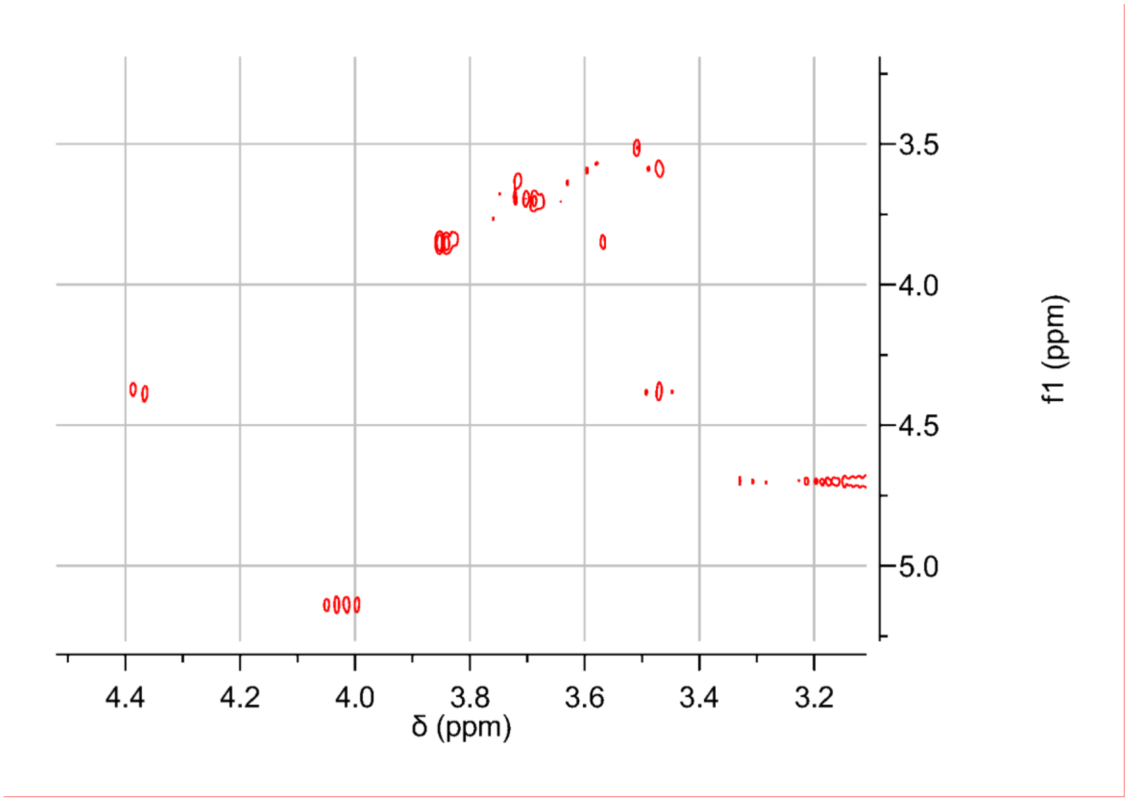


**Fig. S12.**

2D $^1H$-$^1H$ COSY spectrum of larch wood arabinogalactan obtained from abcr GmbH (abcr-LAG).

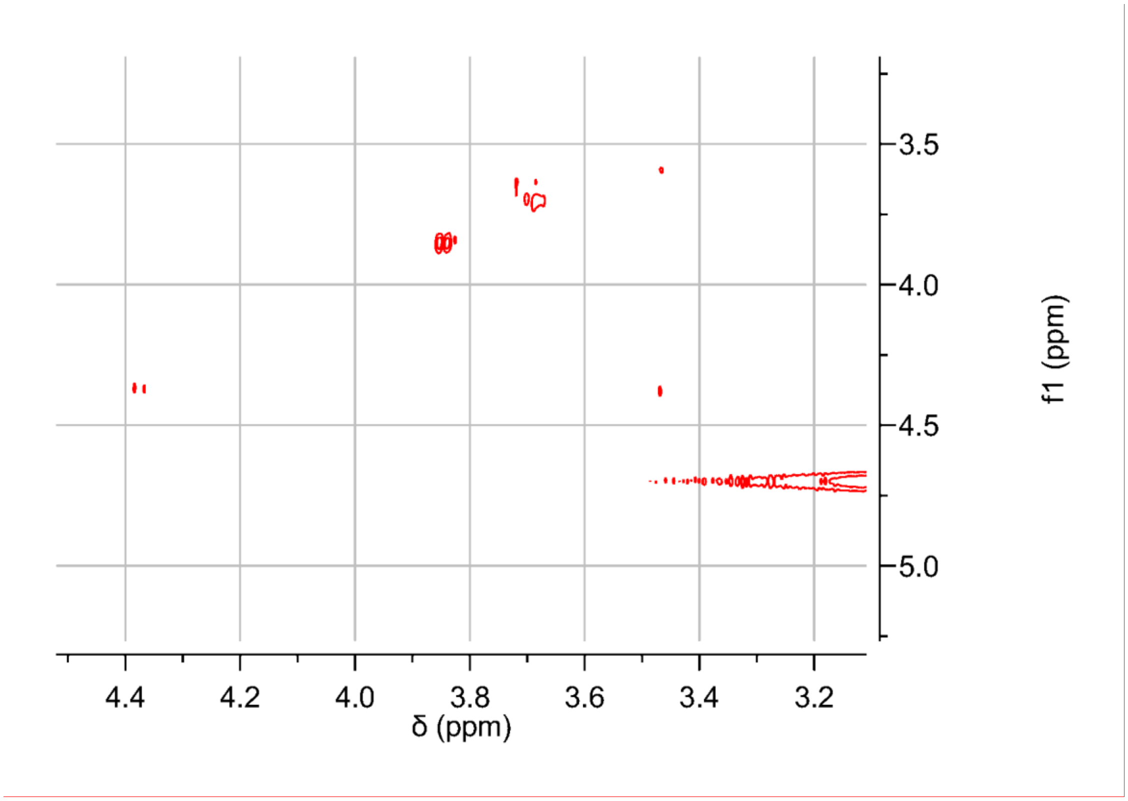


**Fig. S13.**

2D $^{1}H$-$^{1}H$ COSY spectrum of larch wood arabinogalactan obtained from Sigma-Aldrich (Sigma-LAG).

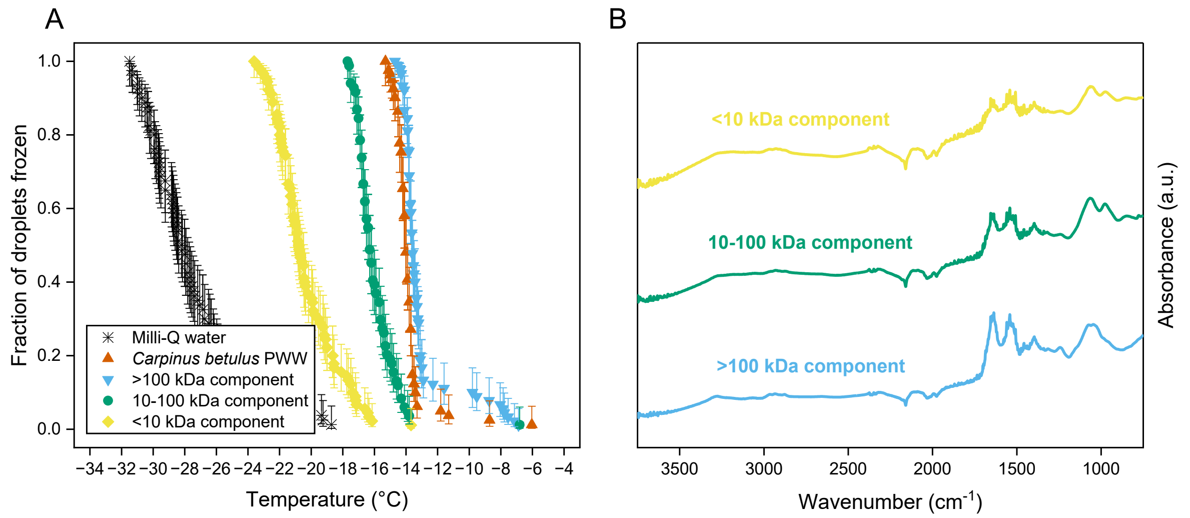


**Fig. S14.**

*Carpinus betulus* pollen washing water (PWW) size separation results: (**A**) Fraction of 1 µL droplets frozen against temperature for Milli-Q® water (background), *Carpinus betulus* PWW and size separated fractions of *Carpinus betulus* PWW: >100 kDa, 10-100 kDa and <10 kDa components. (**B**) Fourier transform infrared absorption spectra for water-soluble material from *Carpinus betulus* pollen >100 kDa, 10-100 kDa and <10 kDa size fractions, fingerprint region only, normalised and offset along the y-axis.

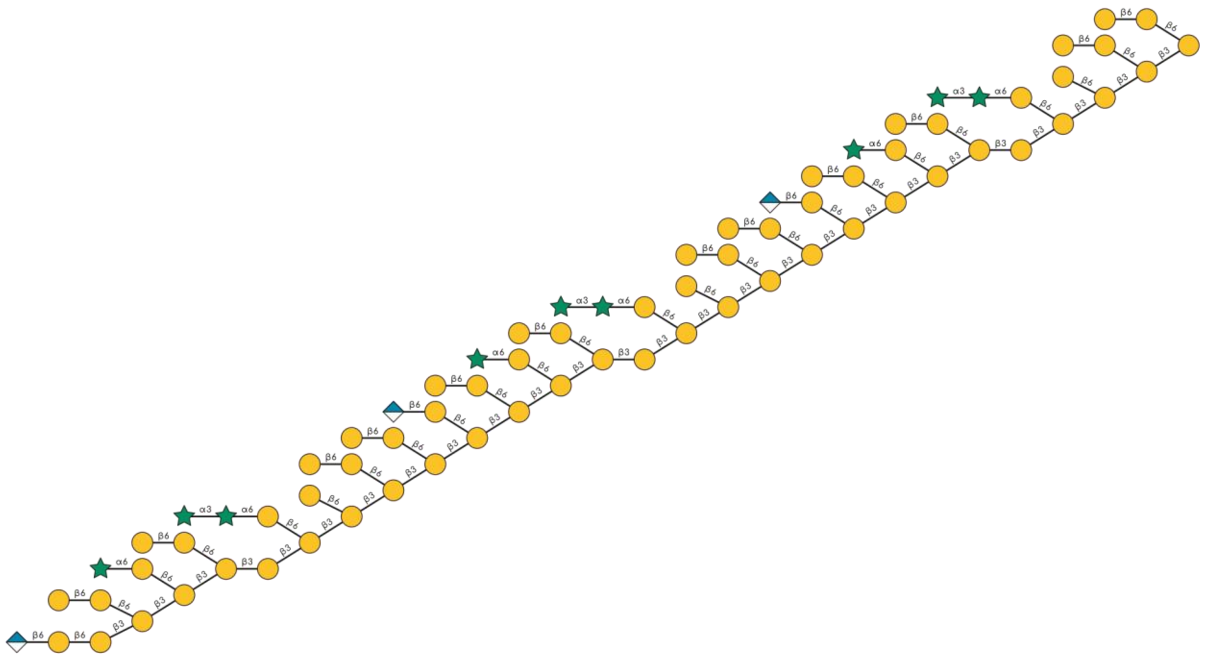

**Fig. S15.**

Symbol Nomenclature for Glycans (SNFG) representation of the branched polysaccharide used in this work. Gal, Ara and GlcA sugars are represented by yellow circles, green stars and white/blue diamonds, respectively.

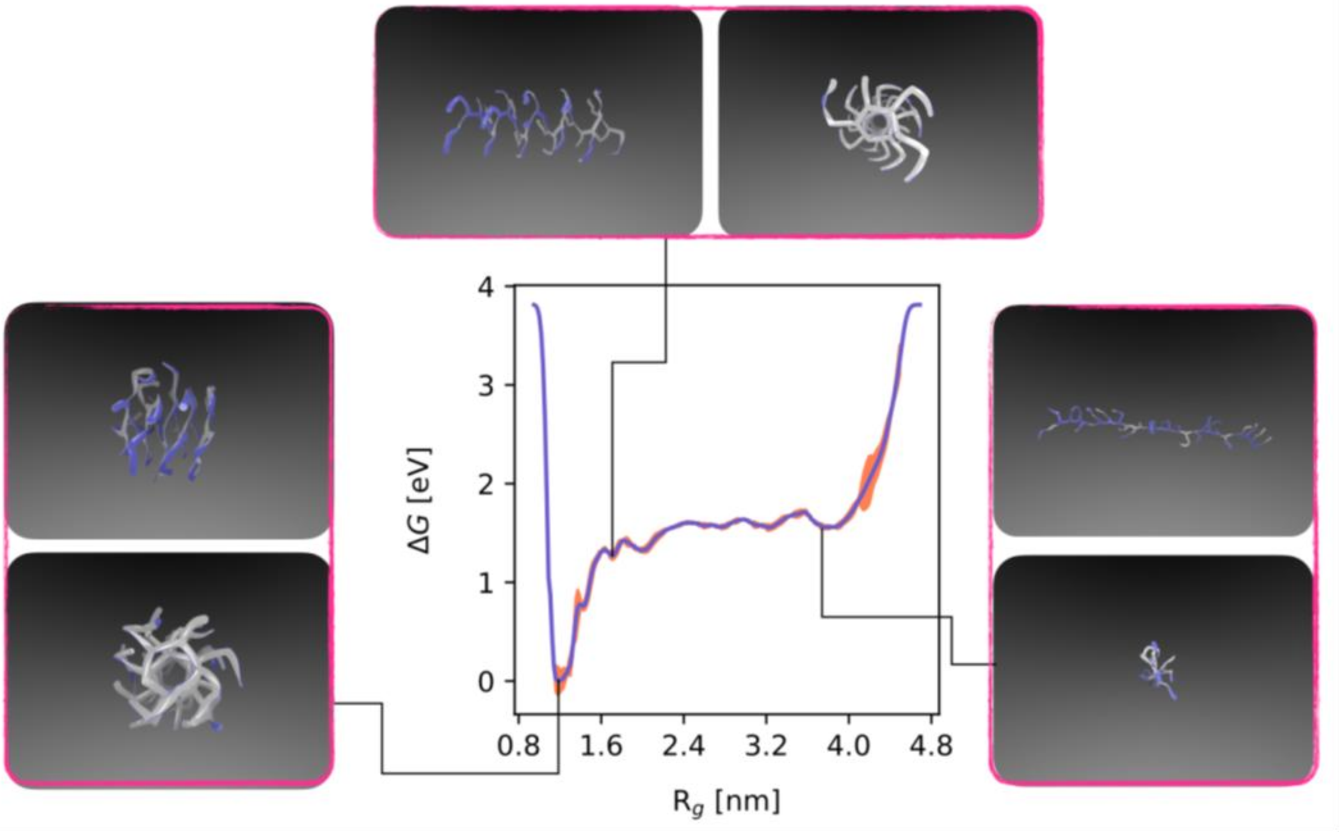


**Fig. S16.**

The conformational space of a 75-sugar LAG model. The free energy profile ($\Delta$G) with respect to the radius of gyration ($R_g$) indicates that the most stable conformation ($R_g$=~1.2 nm) for this system is a helical structure with a shorter pitch re: the structure put forward by CHARMM-GUI, see text ($R_g$=~1.8 nm). As expected, a linear configuration ($R_g$=~3.8 nm) is extremely unlikely to be observed. The orange region illustrates the uncertainty associated with our estimate of $\Delta$G, see text.

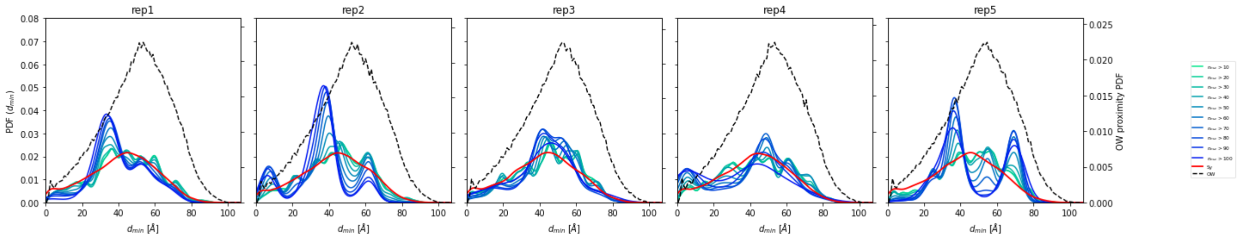


**Fig. S17.**

Spatial distribution of pre-critical ice-like clusters around LAG. Distribution of pre-critical ice-like clusters as a function of distance from the LAG molecule for five statistically independent molecular-dynamics replicas. The synthetic reference distribution is shown in red. A modest enrichment at the LAG–water interface was observed in replicas 2 and 4, but this behaviour was not reproduced consistently across the five replicas.

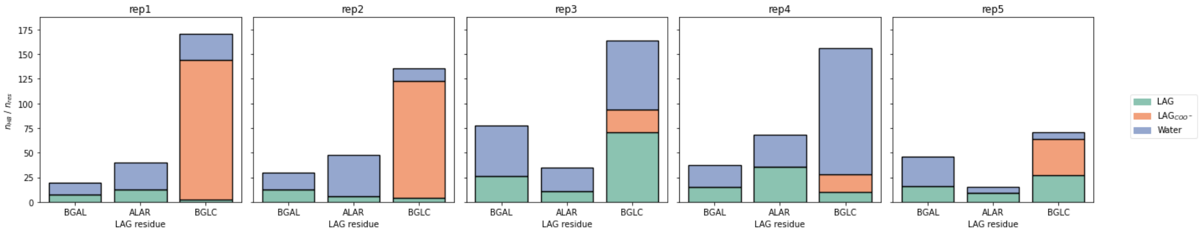


**Fig. S18.**

Contributions of LAG residues to hydrogen bonding with ice-like clusters. Relative involvement of individual LAG residue types in hydrogen bonds formed between the simulated LAG molecule and pre-critical ice-like clusters across five statistically independent replicas. BGLC residues were disproportionately represented despite their low abundance in the model structure.

**Table S1.**

| Species | Collection Date | Plant Location | $T_{50}$ |
|---|---|---|---|
| *Anemia phyllitidis* (L.) Sw. | 17/01/2019 | Kew Gardens | –21.7 °C |
| *Athyrium filix-femina* (L.) Roth | 20/07/2018 | Wakehurst | –12.2 °C |
| *Blechnum spicant* (L.) Roth | 20/07/2018 | Wakehurst | –14.5 °C |
| *Cosentinia vellea* (Aiton) Tod. | 2004 | Sagunto, Valencia, Spain | –19.0 °C |
| *Dicksonia fibrosa* Colenso | 23/01/2019 | Kew Gardens | –21.7 °C |
| *Dryopteris dilatata* (Hoffm.) A.Gray | 18/06/2018 | Wakehurst | –8.7 °C |
| *Dryopteris filix-mas* (L.) Schott | 20/07/2018 | Wakehurst | –15.0 °C |
| *Equisetum arvense* L. | 24/04/2018 | Kew Gardens | –15.5 °C |
| *Equisetum arvense* L. | 06/2019 | -- | –22.6 °C |
| *Hemionitis marantae* (L.) Christenh. | 2003 | Alucudia de Veo, Castellon, Spain | –20.0 °C |
| *Lepisorus fortuni* (T.Moore) C.M.Kuo | 05/2019 | Kew Gardens | –22.5 °C |
| *Onoclea struthiopteris* (L.) Roth | 19/01/2017 | Wakehurst | –14.2 °C |
| *Onoclea struthiopteris* (L.) Roth | 16/02/2018 | Wakehurst | –14.2 °C |
| *Onoclea struthiopteris* (L.) Roth | 01/2019 | Wakehurst | –7.5 °C |
| *Onoclea struthiopteris* (L.) Roth | 27/01/2022 | Wakehurst | –14.1 °C |
| *Onoclea struthiopteris* (L.) Roth | 31/03/2022 | Wakehurst | –16.4 °C |
| *Onoclea struthiopteris* (L.) Roth | 19/05/2022 | Wakehurst | –9.8 °C |
| *Osmundastrum cinnamomeum* (L.) C.Presl | 24/05/2017 | Wakehurst | –18.9 °C |
| *Osmundastrum cinnamomeum* (L.) C.Presl | 25/05/2018 | Wakehurst | –19.0 °C |
| *Osmundastrum cinnamomeum* (L.) C.Presl | 19/05/2022 | Wakehurst | –15.7 °C |
| *Osmunda claytoniana* L. | 25/05/2017 | Wakehurst | –12.9 °C |
| *Osmunda claytoniana* L. | 25/05/2018 | Wakehurst | –20.5 °C |
| *Osmunda regalis* L. | 05/2019 | Wakehurst | –18.5 °C |
| *Osmunda regalis* L. | Not recorded | Wakehurst | –21.9 °C |
| *Microsorum scolopendria* (Burm.f.) Copel. | 2019 | Not recorded | –15.9 °C |
| *Polystichum lonchitis* (L.) Roth | 2003 | Espot, Lleida, Spain | –20.5 °C |
| *Pteris vittata* L. | 2004 or 2005 | Villalonga, Valencia, Spain | –21.5 °C |

Details of fern species, dates and locations of spore collections (where recorded) and mean freezing temperatures, $T_{50}$, of 1 µL droplets of filtered 2 wt% spore washing water (SWW) solutions. Accepted species names were assigned according to the Plants of the World Online Database (*83*). Wakehurst is located in Ardingly, West Sussex, UK and Kew Gardens in Richmond, London, UK (*84*).

**Table S2.**

| Species | Preparation | Collection Date | Plant Location | $T_{50}$ |
|---|---|---|---|---|
| *Funaria hygrometrica* Hedw. | 2 wt% SWW | 24/07/2022 | Wakehurst | –19.8 °C |
| *Polytrichum formosum* Hedw. | 2 wt% SWW | 25/07/2022 | Wakehurst | –17.1°C |
| *Ceratodon purpureus* (Hedw.) Brid. | 2 wt% SWW | 09/08/2025 | Oakley | –17.3 °C |

Details of moss species, dates and locations of collections and mean freezing temperatures, $T_{50}$, of 1 µL droplets. Accepted species names were assigned according to the Bryophyte Nomenclator (*85*) using data from the Tropicos database (*84*). Wakehurst is in Ardingly, West Sussex, UK and Oakley in Bedfordshire, UK.

#### References


1. T. Koop, B. J. Murray, A physically constrained classical description of the homogeneous nucleation of ice in water. *J. Chem. Phys.* **145**, 211915 (2016).
2. D. A. Knopf, P. A. Alpert, Atmospheric ice nucleation. *Nat. Rev. Phys.* **5**, 203–217 (2023).
3. A. Korolev *et al.*, Mixed-phase clouds: Progress and challenges. *Meteorol. Monogr.* **58**, 5.1–5.50 (2017).
4. S. M. Burrows *et al.*, Ice-nucleating particles that impact clouds and climate: Observational and modeling research needs. *Rev. Geophys.* **60**, e2021RG000745 (2022).
5. R. J. Herbert *et al.*, Gaps in our understanding of ice-nucleating particle sources exposed by global simulation of the UK Earth System Model. *Atmos. Chem. Phys.* **25**, 291–325 (2025).
6. M. Z. Jacobson, D. G. Streets, Influence of future anthropogenic emissions on climate, natural emissions, and air quality. *J. Geophys. Res. Atmos.* **114**, D08118 (2009).
7. S. Werchner *et al.*, When do subpollen particles become relevant for ice nucleation processes in clouds? *J. Geophys. Res. Atmos.* **127**, e2021JD036340 (2022).
8. K. Diehl, S. Matthias-Maser, R. Jaenicke, S. K. Mitra, The ice nucleating ability of pollen: Part II. Laboratory studies in immersion and contact freezing modes. *Atmos. Res.* **61**, 125–133 (2002).
9. N. von Blohn, S. K. Mitra, K. Diehl, S. Borrmann, The ice nucleating ability of pollen: Part III. New laboratory studies in immersion and contact freezing modes including more pollen types. *Atmos. Res.* **78**, 182–189 (2005).
10. B. G. Pummer, H. Bauer, J. Bernardi, S. Bleicher, H. Grothe, Suspendable macromolecules are responsible for ice nucleation activity of birch and conifer pollen. *Atmos. Chem. Phys.* **12**, 2541–2550 (2012).
11. S. Augustin *et al.*, Immersion freezing of birch pollen washing water. *Atmos. Chem. Phys.* **13**, 10989–11003 (2013).
12. E. Gute, J. P. D. Abbatt, Oxidative processing lowers the ice nucleation activity of birch and alder pollen. *Geophys. Res. Lett.* **45**, 1647–1653 (2018).
13. E. Gute, J. P. D. Abbatt, Ice nucleating behavior of different tree pollen in the immersion mode. *Atmos. Environ.* **231**, 117488 (2020).
14. J. Burkart, J. Gratzl, T. M. Seifried, P. Bieber, H. Grothe, Isolation of subpollen particles (SPPs) of birch: SPPs are potential carriers of ice nucleating macromolecules. *Biogeosciences* **18**, 5751–5765 (2021).
15. N. L. H. Kinney, C. A. Hepburn, M. I. Gibson, D. Ballesteros, T. F. Whale, High interspecific variability in ice nucleation activity suggests pollen ice nucleators are incidental. *Biogeosciences* **21**, 3201–3214 (2024).
16. B. H. Matthews, A. N. Alsante, S. D. Brooks, Pollen emissions of subpollen particles and ice nucleating particles. *ACS Earth Space Chem.* **7**, 1207–1218 (2023).
17. K. Dreischmeier, C. Budke, L. Wiehemeier, T. Kottke, T. Koop, Boreal pollen contain ice-nucleating as well as ice-binding "antifreeze" polysaccharides. *Sci. Rep.* **7**, 41890 (2017).
18. J. Kretzschmar *et al.*, From trees to rain: Enhancement of cloud glaciation and precipitation by pollen. *Environ. Res. Lett.* **19**, 104052 (2024).

19. Y. Zhang *et al.*, Effects of pollen on hydrometeors and precipitation in a convective system. *J. Geophys. Res. Atmos.* **129**, e2023JD039891 (2024).
20. M. C. Wozniak, F. Solmon, A. L. Steiner, Pollen rupture and its impact on precipitation in clean continental conditions. *Geophys. Res. Lett.* **45**, 7156–7164 (2018).
21. M. Prank, J. Tonttila, X. Shang, S. Romakkaniemi, T. Raatikainen, Can pollen affect precipitation? *Atmos. Chem. Phys.* **25**, 183–197 (2025).
22. P. Wolber, G. Warren, Bacterial ice-nucleation proteins. *Trends Biochem. Sci.* **14**, 179–182 (1989).
23. J. Fröhlich-Nowoisky *et al.*, Ice nucleation activity in the widespread soil fungus *Mortierella alpina*. *Biogeosciences* **12**, 1057–1071 (2015).
24. H. J. Tong *et al.*, A new electrodynamic balance (EDB) design for low-temperature studies: Application to immersion freezing of pollen extract bioaerosols. *Atmos. Meas. Tech.* **8**, 1183–1195 (2015).
25. L. Felgitsch *et al.*, Birch leaves and branches as a source of ice-nucleating macromolecules. *Atmos. Chem. Phys.* **18**, 16063–16079 (2018).
26. B. J. Harris *et al.*, Divergent evolutionary trajectories of bryophytes and tracheophytes from a complex common ancestor of land plants. *Nat. Ecol. Evol.* **6**, 1634–1643 (2022).
27. P. K. Strother, C. Foster, A fossil record of land plant origins from charophyte algae. *Science* **373**, 792–796 (2021).
28. S. Wallace, A. Fleming, C. H. Wellman, D. J. Beerling, Evolutionary development of the plant spore and pollen wall. *AoB Plants* **2011**, plr027 (2011).
29. T. F. Whale, Quantification of the ice nucleation activity of ice-binding proteins using a microliter droplet freezing experiment. *Methods Mol. Biol.* **2730**, 121–134 (2024).
30. B. G. Pummer *et al.*, Ice nucleation by water-soluble macromolecules. *Atmos. Chem. Phys.* **15**, 4077–4091 (2015).
31. M. Kačuráková, P. Capek, V. Sasinková, N. Wellner, A. Ebringerová, FT-IR study of plant cell wall model compounds: Pectic polysaccharides and hemicelluloses. *Carbohydr. Polym.* **43**, 195–203 (2000).
32. M. I. Daily, M. D. Tarn, T. F. Whale, B. J. Murray, An evaluation of the heat test for the ice-nucleating ability of minerals and biological material. *Atmos. Meas. Tech.* **15**, 2635–2665 (2022).
33. J. L. Morris *et al.*, The timescale of early land plant evolution. *Proc. Natl. Acad. Sci. U.S.A.* **115**, E2274–E2283 (2018).
34. P. C. J. Donoghue, C. J. Harrison, J. Paps, H. Schneider, The evolutionary emergence of land plants. *Curr. Biol.* **31**, R1281–R1298 (2021).
35. X. Ma, C. Zhang, L. Yang, S. B. Hedges, B. Zhong, New insights on angiosperm crown age based on Bayesian node dating and skyline fossilized birth-death approaches. *Nat. Commun.* **16**, 2265 (2025).
36. B. Classen, A. Baumann, J. Utermoehlen, Arabinogalactan-proteins in spore-producing land plants. *Carbohydr. Polym.* **210**, 215–224 (2019).
37. T. M. Seifried *et al.*, Fluorescence signal of proteins in birch pollen distorted within its native matrix: Identification of the fluorescence suppressor quercetin-3-*O*-sophoroside. *Anal. Bioanal. Chem.* **414**, 7531–7542 (2022).
38. F. Wieland *et al.*, Aggregation of ice-nucleating macromolecules from *Betula pendula* pollen determines ice nucleation efficiency. *Biogeosciences* **22**, 103–115 (2025).

39. D. Bartels *et al.*, Evolution of plant cell wall: Arabinogalactan-proteins from three moss genera show structural differences compared to seed plants. *Carbohydr. Polym.* **163**, 227–235 (2017).
40. A. Clarke, R. Anderson, B. Stone, Form and function of arabinogalactans and arabinogalactan-proteins. *Phytochemistry* **18**, 521–540 (1979).
41. A. Majewska-Sawka, E. A. Nothnagel, The multiple roles of arabinogalactan proteins in plant development. *Plant Physiol.* **122**, 3–10 (2000).
42. A. M. Showalter, Arabinogalactan-proteins: Structure, expression and function. *Cell. Mol. Life Sci.* **58**, 1399–1417 (2001).
43. Y. Ma, K. Johnson, Arabinogalactan proteins: Multifunctional glycoproteins of the plant cell wall. *Cell Surf.* **9**, 100102 (2023).
44. K.-K. Mueller, L. Pfeifer, L. Wegner, K. Ehlers, B. Classen, New insights into bryophyte arabinogalactan-proteins from a hornwort and a moss model organism. *Plant J.* **123**, e70312 (2025).
45. T. A. Torode *et al.*, Branched pectic galactan in phloem-sieve-element cell walls: Implications for cell mechanics. *Plant Physiol.* **176**, 1547–1558 (2018).
46. T. L. Lowary, Twenty years of mycobacterial glycans: Furanosides and beyond. *Acc. Chem. Res.* **49**, 1379–1388 (2016).
47. K. Kitazawa *et al.*, β-Galactosyl Yariv reagent binds to the β-1,3-galactan of arabinogalactan proteins. *Plant Physiol.* **161**, 1117–1126 (2013).
48. T. F. Whale, T. Surawy-Stepney, S. L. Barr, Survival-analysis methods for droplet-freezing data: Kaplan–Meier confidence intervals and non-parametric curve comparison. *Atmos. Meas. Tech.* **19**, 6699–6718 (2026).
49. N. H. Fletcher, Size effect in heterogeneous nucleation. *J. Chem. Phys.* **29**, 572–576 (1958).
50. V. A. Babkin, N. A. Neverova, E. N. Medvedeva, T. E. Fedorova, A. A. Levchuk, Investigation of physicochemical properties of arabinogalactan of different larch species. *Russ. J. Bioorg. Chem.* **42**, 707–711 (2016).
51. V. R. Després *et al.*, Primary biological aerosol particles in the atmosphere: A review. *Tellus B Chem. Phys. Meteorol.* **64**, 15598 (2012).
52. B. van Zanten, Experimental studies on trans-oceanic long-range dispersal of moss spores in the Southern Hemisphere. *J. Hattori Bot. Lab.* **44**, 455–482 (1978).
53. C. Hervé *et al.*, Arabinogalactan proteins have deep roots in eukaryotes: Identification of genes and epitopes in brown algae and their role in *Fucus serratus* embryo development. *New Phytol.* **209**, 1428–1441 (2016).
54. T. F. Whale *et al.*, A technique for quantifying heterogeneous ice nucleation in microlitre supercooled water droplets. *Atmos. Meas. Tech.* **8**, 2437–2447 (2015).
55. C. I. Biggs *et al.*, Mimicking the ice recrystallization activity of biological antifreezes: When is a new polymer "active"? *Macromol. Biosci.* **19**, 1900082 (2019).
56. D. M. Murphy, T. Koop, Review of the vapour pressures of ice and supercooled water for atmospheric applications. *Q. J. R. Meteorol. Soc.* **131**, 1539–1565 (2005).
57. D. Barahona, Analysis of the effect of water activity on ice formation using a new thermodynamic framework. *Atmos. Chem. Phys.* **14**, 7665–7680 (2014).
58. E. M. Goellner, J. Utermoehlen, R. Kramer, B. Classen, Structure of arabinogalactan from *Larix laricina* and its reactivity with antibodies directed against type-II arabinogalactans. *Carbohydr. Polym.* **86**, 1739–1744 (2011).

59. S.-J. Park *et al.*, CHARMM-GUI Glycan Modeler for modeling and simulation of carbohydrates and glycoconjugates. *Glycobiology* **29**, 320–331 (2019).
60. O. Guvench *et al.*, CHARMM additive all-atom force field for carbohydrate derivatives and its utility in polysaccharide and carbohydrate–protein modeling. *J. Chem. Theory Comput.* **7**, 3162–3180 (2011).
61. J. L. F. Abascal, E. Sanz, R. García Fernández, C. Vega, A potential model for the study of ices and amorphous water: TIP4P/Ice. *J. Chem. Phys.* **122**, 234511 (2005).
62. C. M. Miles, P.-C. Hsu, A. M. Dixon, S. Khalid, G. C. Sosso, Lipid bilayers as potential ice nucleating agents. *Phys. Chem. Chem. Phys.* **24**, 6476–6491 (2022).
63. R. Chandrasekaran, S. Janaswamy, Morphology of Western larch arabinogalactan. *Carbohydr. Res.* **337**, 2211–2222 (2002).
64. A. Barducci, G. Bussi, M. Parrinello, Well-tempered metadynamics: A smoothly converging and tunable free-energy method. *Phys. Rev. Lett.* **100**, 020603 (2008).
65. M. Bonomi *et al.*, Promoting transparency and reproducibility in enhanced molecular simulations. *Nat. Methods* **16**, 670–673 (2019).
66. M. Salvalaglio, P. Tiwary, M. Parrinello, Assessing the reliability of the dynamics reconstructed from metadynamics. *J. Chem. Theory Comput.* **10**, 1420–1425 (2014).
67. D. van der Spoel *et al.*, GROMACS: Fast, flexible, and free. *J. Comput. Chem.* **26**, 1701–1718 (2005).
68. M. A. Cuendet, W. F. van Gunsteren, On the calculation of velocity-dependent properties in molecular dynamics simulations using the leapfrog integration algorithm. *J. Chem. Phys.* **127**, 184102 (2007).
69. S. Miyamoto, P. A. Kollman, SETTLE: An analytical version of the SHAKE and RATTLE algorithm for rigid water models. *J. Comput. Chem.* **13**, 952–962 (1992).
70. B. Hess, P-LINCS: A parallel linear constraint solver for molecular simulation. *J. Chem. Theory Comput.* **4**, 116–122 (2008).
71. G. Bussi, D. Donadio, M. Parrinello, Canonical sampling through velocity rescaling. *J. Chem. Phys.* **126**, 014101 (2007).
72. M. Bernetti, G. Bussi, Pressure control using stochastic cell rescaling. *J. Chem. Phys.* **153**, 114107 (2020).
73. G. A. Tribello, F. Giberti, G. C. Sosso, M. Salvalaglio, M. Parrinello, Analyzing and driving cluster formation in atomistic simulations. *J. Chem. Theory Comput.* **13**, 1317–1327 (2017).
74. P. J. Steinhardt, D. R. Nelson, M. Ronchetti, Bond-orientational order in liquids and glasses. *Phys. Rev. B* **28**, 784–805 (1983).
75. P. Rodger, T. Forester, W. Smith, Simulations of the methane hydrate/methane gas interface near hydrate-forming conditions. *Fluid Phase Equilib.* **116**, 326–332 (1996).
76. F. Bachtiger, T. R. Congdon, C. Stubbs, M. I. Gibson, G. C. Sosso, The atomistic details of the ice recrystallisation inhibition activity of PVA. *Nat. Commun.* **12**, 1323 (2021).
77. R. J. Gowers *et al.*, "MDAnalysis: A Python package for the rapid analysis of molecular dynamics simulations," in *Proceedings of the 15th Python in Science Conference*, S. Benthall, S. Rostrup, Eds. (SciPy, Austin, TX, 2016), pp. 98–105.
78. C. P. Garnham, R. L. Campbell, P. L. Davies, Anchored clathrate waters bind antifreeze proteins to ice. *Proc. Natl. Acad. Sci. U.S.A.* **108**, 7363–7367 (2011).

79. C. F. Weber, *Polytrichum commune* spores nucleate ice and associated microorganisms increase the temperature of ice nucleation activity onset. *Aerobiologia* **32**, 353–361 (2016).
80. B. F. Moffett, Ice nucleation in mosses and liverworts. *Lindbergia* **38**, 14–16 (2015).
81. J. Yariv, H. Lis, E. Katchalski, Precipitation of arabic acid and some seed polysaccharides by glycosylphenylazo dyes. *Biochem. J.* **105**, 1C–2C (1967).
82. M. Jermyn, Y. Yeow, E. Woods, A class of lectins present in the tissues of seed plants. *Aust. J. Plant Physiol.* **2**, 501–531 (1975).
83. A. Clarke, P. Gleeson, M. Jermyn, R. Knox, Characterization and localization of β-lectins in lower and higher plants. *Aust. J. Plant Physiol.* **5**, 707–722 (1978).
84. POWO, *Plants of the World Online* (Royal Botanic Gardens, Kew, 2026); https://powo.science.kew.org/ (accessed 5 November 2026).
85. Tropicos.org, Missouri Botanical Garden, St. Louis, MO; https://www.tropicos.org/ (accessed 5 November 2026).
86. J.C. Brinda, J.J. Atwood, Eds., *The Bryophyte Nomenclator*; https://www.bryonames.org/ (accessed 5 November 2026).
